# Path sampling simulations reveal how the Q61L mutation restricts the dynamics of KRas

**DOI:** 10.1101/2020.02.28.969451

**Authors:** Sander Roet, Ferry Hooft, Peter G. Bolhuis, David W.H. Swenson, Jocelyne Vreede

## Abstract

The GTPase KRas is a signaling protein in networks for cell differentiation, growth, and division. KRas mutations can prolong activation of these networks, resulting in tumor formation. When active, KRas tightly binds GTP. Several oncogenic mutations affect the conversion between this rigid state and inactive, more flexible states. Detailed understanding of these transitions may provide valuable insights into how mutations affect KRas. Path sampling simulations, which focus on transitions, show KRas visiting several states, which are the same for wild type and the oncogenic mutant Q61L. Large differences occur when converting between these states, indicating the dramatic effect of the Q61L mutation on KRas dynamics. For Q61L a route to the flexible state is inaccessible, thus shifting the equilibrium to more rigid states. Our methodology presents a novel way to predict dynamical effects of KRas mutations, which may aid in identifying potential therapeutic targets.

**Author summary:** Cancer cells frequently contain mutations in the protein KRas. However, KRas is a challenging target for anti-cancer drugs, in part because the dynamic behavior of flexible regions in the protein is difficult to characterize experimentally, and occurs on timescales that are too long for straightforward molecular dynamics simulations. We have used path sampling, an advanced simulation technique that overcomes long timescales, to obtain atomistic insight into the dynamics of KRas. Comparing the oncogenic mutant Q61L to the wild type revealed that the mutation closes off one transition channel for deactivating KRas. Our approach opens up the way for predicting the dynamical effects of mutations in KRas, which may aid in identifying potential therapeutic targets.

## Introduction

Ras GTPases are signal transduction proteins that mediate cell growth, cell differentiation and death, and comprise the most frequently occurring family of oncoproteins in human cancers [1, 2]. Mutations in Ras proteins initiate cell transformation, drive oncogenesis and promote tumor maintenance. The Ras family of oncoproteins has been studied extensively for almost three decades, as activation of Ras represents a key feature of malignant transformation for many cancers. In the cancers that contribute most heavily to worldwide mortality, Ras mutations are extremely common [3]. Many Ras-activating mutations have been detected in non-small cell lung cancer (15 to 20% of tumors), colon adenomas (40%) and pancreatic adenocarcinomas (95%) making it the single most common human oncoprotein. Binding of guanosine triphosphate (GTP) activates signal transduction by Ras proteins, while their GTPase function inactivates signal transduction again by hydrolyzing GTP to guanosine diphosphate (GDP). Ras proteins consist of a highly conserved catalytic domain called the G domain and a variable C domain which anchors Ras in the membrane. Several isoforms exist of Ras, which are implicated in different types of cancer [2, 3]. A member of this family, KRas-4B, is often found in common and life-threatening cancers, such as lung cancer, colon cancer and pancreatic cancer [3].

In this work we focus on the G domain of the KRas-4B isoform, which contains 166 residues and can be considered as the minimal signaling unit. This domain contains the guanine nucleotide binding site and two regions that sense the nature of the bound nucleotide, switch 1, S1, and switch 2, S2. Literature has not reached consensus on which range of residues correspond to each switch region [4, 5]. We chose to use a narrow definition that corresponds to the residues that are important for the conformational changes in this study, using residues 30-33 for S1 and residues 60–66 for S2. These regions, highlighted in green (S1) and blue (S2) in figure 1(left), are involved in many interactions between Ras and partners. In the GTP-bound state, Ras interacts with downstream effectors such as the Raf and PI3K kinases [6]. After hydrolysis of GTP, these loop regions adopt a more open conformation [7] and exhibit more flexibility, causing Ras to lose the ability to bind to downstream effectors. While bound to GTP, Ras exists in a dynamic equilibrium between a weakly populated state 1 and a dominant state 2 [8, 9]. Conformational state 1 is more flexible and open than the closed, ordered state 2, as schematically shown in figure 1(right). Crystal structures of Ras bound with GTP analogues are typically in the state 2 form of Ras [6, 10]. However, ^31^P-NMR studies report that the switch regions can also adopt disordered conformations when bound to GTP, similar to the GDP-bound state [11]. This work will focus on the transition between the ordered (state 2) and disordered (state 1) states of GTP-bound KRas-4B.

**Fig 1.**
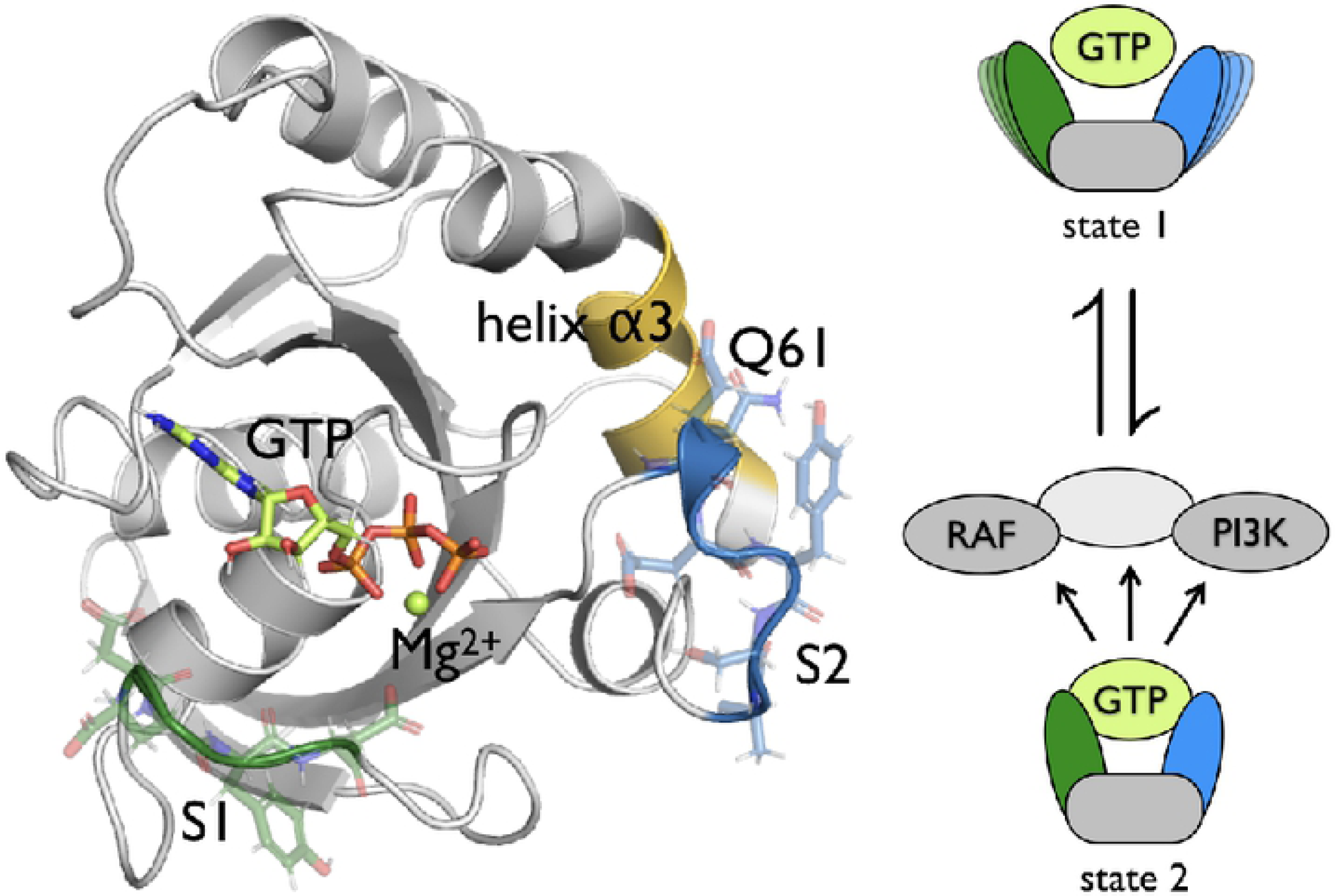
Ras structure and function. Structure of GTP-bound KRas in the active state 2 (left). The switch regions are highlighted in green for S1 and blue for S2, the *α*3-helix is highlighted in yellow. The protein is shown as a ribbon with an transparent stick representation for the amino acids in S1 and S2. GTP is shown as solid sticks, with carbon atoms colored in green, oxygen in red, nitrogen in blue and phosphorus in orange. Mg^2+^ is shown as a green ball. Schematic representation of the inactive state 1 and the active state 2 of GTP-bound KRas (right). The S1 region is represented in green, the S2 region in blue, and the rest of the protein in grey. State 1 corresponds to the conformational state in which S1 and S2 are more flexible and not bound to GTP. State 2 corresponds to the conformational state in which both S1 and S2 are bound to GTP. State 2 activates downstream effectors like RAF and PI3K by binding them.

Ras activating mutations include substitutions at glycine (G) 12, G13 and glutamine (Q) 61 [12], which affect the conformational equilibrium of Ras. Replacing Q61, located in S2, results in reduced GTPase activity in Ras [13] and an altered conformational space for KRas-4B [14]. Changing Q61 into leucine (L) results in an oncogenic mutant [4]. By replacing a hydrophilic glutamine with a hydrophobic leucine [15], the conformational space of GTP-bound KRas-4B will change, and alter the transition between state 1 and state 2, causing the Q61L mutant to be more ordered. As a consequence, KRas-4B will remain more in an activating role, thus promoting cell growth and cell differentiation, and eventually tumor formation.

As the Q61L mutation may affect the conformations of state 1 and state 2, as well as the transitions between these states, methods that provide high resolution in both space and time are required. Molecular simulations can provide such high detail. Previous molecular simulation studies of Ras proteins focused on the dynamics of the stable states and the effect of oncogenic mutations on those dynamics, as described in the overview by Prakash et al. [16]. It is still an open question how Ras converts from ordered to less ordered conformational states. In particular the effect of mutations on these transitions is unclear.

Currently, a brute-force all-atom molecular dynamics (MD) investigation of the mechanism and kinetic aspects of the conformational transitions in Ras is not feasible, as the time scales involved are in the order of hundreds of microseconds, [17, 18]. Such long timescales are caused by high free energy barriers separating the stable states. One way of overcoming these barriers is by employing biasing potentials that drive the system towards the barrier region along a predefined reaction coordinate. For HRas, a steered MD study revealed the importance of water molecules in entering the active state 2. [19]. While methods employing additional potentials are well suited for computing free-energy barriers and other thermodynamic properties, they often fail to yield mechanistic insight at ambient conditions, as a poor choice of reaction coordinate may lead to a wrong reaction mechanism, bad sampling and a poor estimation of the rate constants.

The transition path sampling (TPS) algorithm [20] is another way to address the timescale problem, which avoids these drawbacks. TPS is a Monte Carlo (MC) simulation in the space of trajectories and collects an ensemble of short reactive trajectories connecting a predefined initial and final state, without prior knowledge of the transition state region. The speed-up gained by using TPS and related techniques is tremendous. Assuming a transition rate in the order of 10 s^−1^, observing a single transition would require on average 100 ms of MD. In contrast, when using TPS, the barrier region is sampled using MD trajectories of only tens of nanoseconds, thus providing a speed up in the order of several million to a billion. Path sampling methods have been extended from the original two-state formalism to be used with multiple stable states [21]. At a given MC step, a TPS simulation samples a transition between a specific pair of states. However, in the multiple state TPS (MSTPS) approach, it is possible to generate trajectories that sample a different pair of states. The frequency of this switching between transitions depends on the barrier separating different transition channels. Analysis of the switching behavior can provide useful insight into the overall dynamics.

For both S1 and S2, three stable states were identified that could be used in MSTPS simulations of the wild type (WT) as well as Q61L. S2 displays different dynamics for WT and Q61L. While the WT simulations frequently switch from one transition to another, the Q61L simulations hardly switch at all. Closer examination reveals that the WT S2 can reach the flexible open state via a channel that is not accessible in Q61L. Both WT and Q61L can reach the open state by S2 sliding along a slightly hydrophobic pocket of the *α*3-helix. However, the Q61L mutation prevents direct solvation of S2, which is possible for the WT protein. This observation may also explain why Q61L occurs more often as an oncogenic mutation in HRas. [4] As the *α*3-helix of HRas is slightly less hydrophilic compared to KRas, it would have a stronger interaction with S2, resulting in fewer transitions into the open state. As a result, the open, inactive state will occur less frequently and the protein is more likely to be in an ordered state, thus prolonging the activation signaling networks, ultimately leading to extended cell growth and division.

Our results show that our methodology is able to map out the dynamics of a Ras protein, can indicate differences in dynamics between a WT protein and an oncogenic mutant, and is able to reveal details on the nature of the altered behavior as caused by the mutation. As such, our approach presents a powerful tool in providing a better understanding of the dynamic nature of Ras proteins, and may potentially aid in identifying new therapeutic targets for tumors induced by Ras mutations.

## Materials and methods

### 1 Methods and Materials

#### Structure generation

The initial GTP-bound KRas-4B structure was constructed from the crystal structure of GppNHp bound HRas (PDB-code: 4EFL) [17, 18]. This was done by first using homology modeling (MODELLER v9.16) [22], using sequential alignment to convert HRas to KRas-4B. Then the GppNHp was manually modified into GTP, by changing the nitrogen into an oxygen and removing the attached hydrogen. Finally, structures of the protein and the GTP were combined into a single file. The initial structure for the mutant (Q61L) was made from this structure by mutating the glutamine (Q) 61 of this final structure into a leucine (L), using MODELLER [22].

The initial structures were put inside a dodecahedral periodic box with a minimum distance between the structures and the side of the box of 1 nm. This resulted in boxes with volumes of 228.154 nm^3^ and 230.723 nm^3^ for the wild type (WT) and Q61L, respectively. The boxes were filled with TIP3P water [23]. 51 of the waters were replaced by 30 Na^+^ and 21 Cl^−^ ions to neutralize the systems and achieve a physiological salt concentration of 0.15 M NaCl. This resulted in total system sizes of 22561 and 22857 atoms for the WT and Q61L, respectively.

#### Molecular Dynamics

##### Procedure

The initial systems were equilibrated in four steps, consisting of energy minimization, an isothermal equilibration, an isothermal-isobaric equilibration and a 1 ns molecular dynamics simulation. The equilibrated structures were used to run four 100 ns molecular dynamics simulations for both WT and Q61L.

##### Settings

In the molecular dynamics simulations the atomic interactions were described by the AMBER99SB-ILDN [24] force field, extended with optimized parameters for the triphosphate chain of GTP [25]. Long-range electrostatic interactions were treated via the Particle Mesh Ewald method [26]. The short-range non-bonded interactions (e.g. electrostatics and Van der Waals interactions) were cut off at 1.1 nm.

All of the equilibration was performed with GROMACS v.4.6.5. [27]. The leap-frog integrator was used with a time step of 2 fs. Temperature was kept constant at 310 K using the v-rescale thermostat [28] using two temperature coupling groups: the first group consisted of the protein, GTP and Mg^2+^, while the second group consisted of water, Na^+^, and Cl^−^. The pressure was kept constant using the Parrinello-Rahman barostat [29] at a pressure of 1 bar. All bond lengths were constrained using the LINCS algorithm [30].

The 100 ns production runs were performed with OpenMM (7.1.0.dev-5e53567) [31]. The constraints were changed to only constrain the bond lengths of bonds to a hydrogen, using SHAKE [32], the integrator was the Velocity Verlet with velocity randomization (VVVR) integrator [33] from OpenMMTools v.0.14 [34] and the barostat was the Monte Carlo barostat [35]. The production simulations were run using the CUDA platform of OpenMM on NVIDIA GeForce GTX TITAN X GPUs.

#### Collective variables and stable states

The long molecular dynamics runs were visually analyzed to identify stable states, using VMD [36]. Five types of collective variable functions were used to define the stable states, which are described in table 2 in Appendix 1.

**Table 1.** Statistics of the S2 MSTPS simulations. MC steps indicates the number of Monte-Carlo trials, also called shooting moves. Accepted steps refers to the number of MC trials that were accepted and the acceptance is 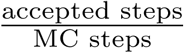. Decorrelated trajectories indicate the number of accepted trajectories that do not have any frames in common. The average path lengths is the sum of the length of the trajectories, weighted by their MC weight, divided by the number of MC steps. The total simulation time is the total time of MD performed by the MD engine in the MSTPS simulations.

|  | WT |  |  | Q61L |  |  |
| --- | --- | --- | --- | --- | --- | --- |
|  | sim 1 | sim2 | sim 3 | sim 1 | sim 2 | sim 3 |
| MC steps | 2000 | 2000 | 2000 | 2000 | 2000 | 2000 |
| Accepted steps | 766 | 824 | 787 | 688 | 748 | 759 |
| Acceptance | 38.30% | 41.20% | 39.35% | 34.40% | 37.38% | 37.95% |
| Decorrelated trajectories | 131 | 151 | 128 | 99 | 129 | 125 |
| Average path length | 3.49 ns | 1.26 ns | 1.50 ns | 1.42 ns | 1.36 ns | 1.98 ns |
| Total simulation time | 6.01 μs | 2.15 μs | 2.10 μs | 2.26 μs | 1.83 μs | 3.74 μs |

**Table 2.** List of the different collective variable types.

| CV type | Description |
| --- | --- |
| Minimum distance | The smallest distance between two groups of atoms. Using MDTraj [43], distances of every atom pair were calculated. The lowest is the minimum distance. |
| Circular mean center of mass (cCOM)* | The circular mean center of mass (cCOM) is a center of geometry calculation that allows for periodicity. The system is first mapped onto a cube, followed by the calculation of the center of mass, using the procedure of ref. [44]. Then the cCOM is mapped back onto the original axes. |
| Number of hydrogen bonds | The number of hydrogen bonds is calculated by counting how many of the possible donor-acceptor pairs form a hydrogen bond. A hydrogen bond in this code is defined by having a $H_{donor}$-acceptor distance smaller than 0.25 nm and having an $X_{donor}$-$H_{donor}$-acceptor angle larger than $\frac{2}{3}\pi$ rad. |
| Number of water mediated hydrogen bonds | The number of water mediated hydrogen bonds is calculated by first selecting all water oxygens that are within a distance of 0.35 nm from both input groups. If a water forms a hydrogen bond with both groups, it is counted as a water mediated hydrogen bond. The hydrogen bond calculation is done as described above. |
| Number of bonds | The number of bonds is the number of pairs, from a given list of pairs, for which the minimum distance is smaller than 0.35 nm. The minimum distance is defined in the minimum distance cv type. |

\* Note that this circular mean center of mass does not give the actual center of mass.
In the CV type column the name of the collective variable type as used in table 3 and 4 are shown, with their description in the Description column.

For both WT and Q61L the relevant collective variables can be found in table 3 and 4, for S1 and S2, respectively. These collective variables are comprised of the collective variable types described in table 2. The stable states for S1 were S1-D33, S1-30-32, and S1-open, and for S2 were S2-GTP, S2-*α*3, and S2-open. The definitions of all stable states can be found in table 5.

**Table 3.**
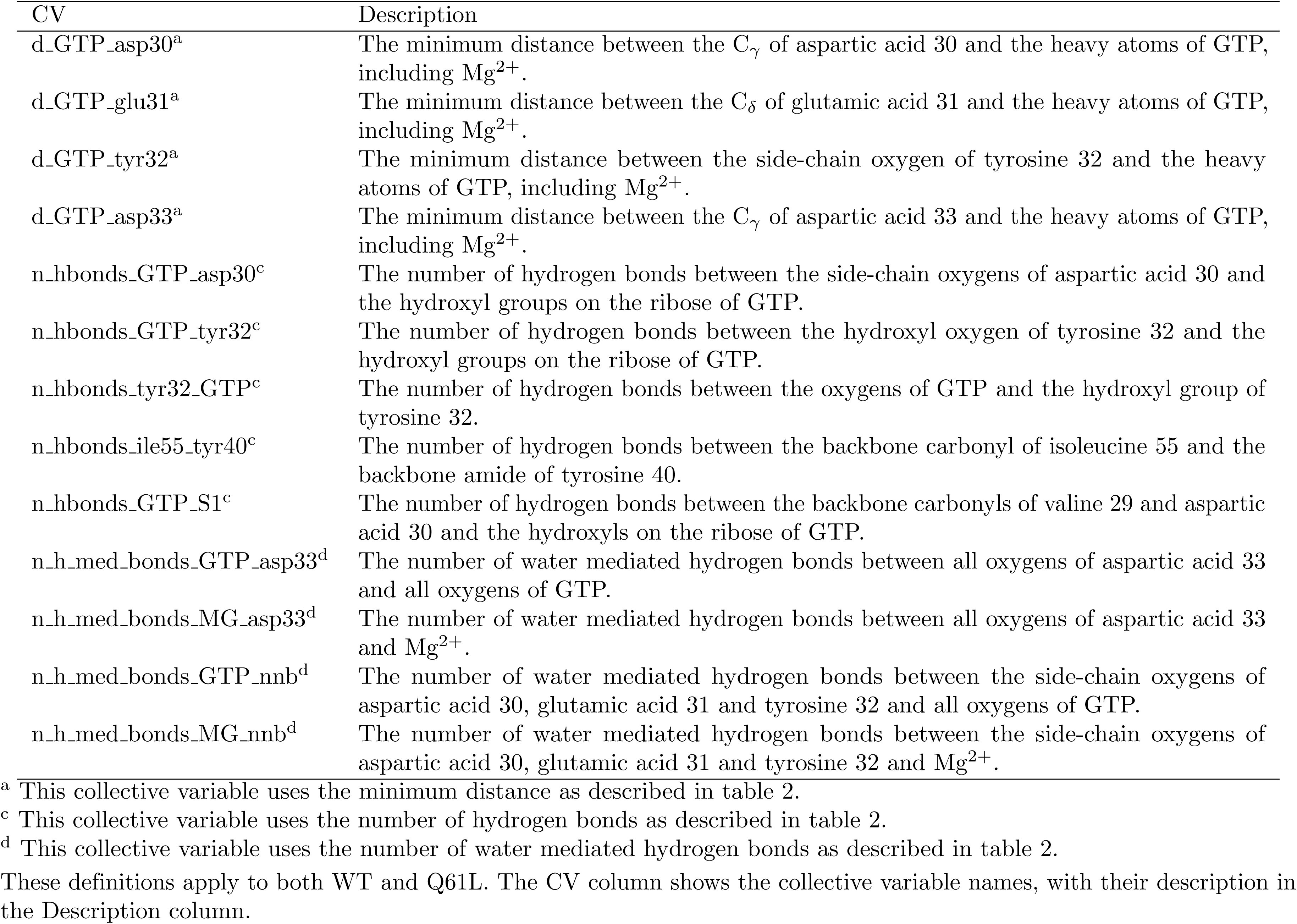
List of the relevant collective variables for the stable state definitions of S1.

**Table 4.** List of the relevant collective variables for the stable state definitions of S2.

| CV | Description |
| --- | --- |
| d_gly12_gly60<sup>a</sup> | The minimum distance between the heavy atoms of glycine 12 and the heavy atoms of glycine 60. |
| d_gly12_gln61<sup>a,wt</sup> | The minimum distance between the heavy atoms of glycine 12 and the side-chain heavy atoms of glutamine 61. |
| d_gly12_leu61<sup>a,Q61L</sup> | The minimum distance between the heavy atoms of glycine 12 and the side-chain heavy atoms of leucine 61. |
| d_GTP_glu62<sup>a</sup> | The minimum distance between the C<sub>δ</sub> of glutamic acid 62 and the heavy atoms of GTP, including Mg<sup>2+</sup>. |
| d_GTP_glu63<sup>a</sup> | The minimum distance between the C<sub>δ</sub> of glutamic acid 63 and the heavy atoms of GTP, including Mg<sup>2+</sup>. |
| d_cCOM_GTP_S2<sup>a,b</sup> | The minimum distance between the circular mean center of mass of all atoms of residues 61 to 66 and the circular mean center of mass of all atoms of GTP, including Mg<sup>2+</sup>. |
| n_S2_α3<sup>e</sup> | The number of combinations between the sets of {histidine 95, tyrosine 96, glutamine 99, arginine 102} and {{61}<sup>*</sup>, glutamic acid 62, glutamic acid 63, tyrosine 64} for which the minimal distance between the side-chain heavy atoms of the residue from the first set and all heavy atoms of the residue from the second set is smaller than 0.35 nm. |

<sup>a</sup> This collective variable uses the minimum distance as described in table 2.
<sup>b</sup> This collective variable uses the circular mean center of mass as described in table 2.
<sup>e</sup> This collective variable uses the number of bonds as described in table 2.
<sup>wt</sup> Only used in wild-type KRas.
<sup>Q61L</sup> Only used in the Q61L mutant of KRas.
<sup>\*</sup> {61} is glutamine 61 for wild-type KRas and leucine 61 for the Q61L mutant of KRas.
These definitions apply to both WT and Q61L. The CV column shows the collective variable names, with their description in the Description column.

**Table 5.**
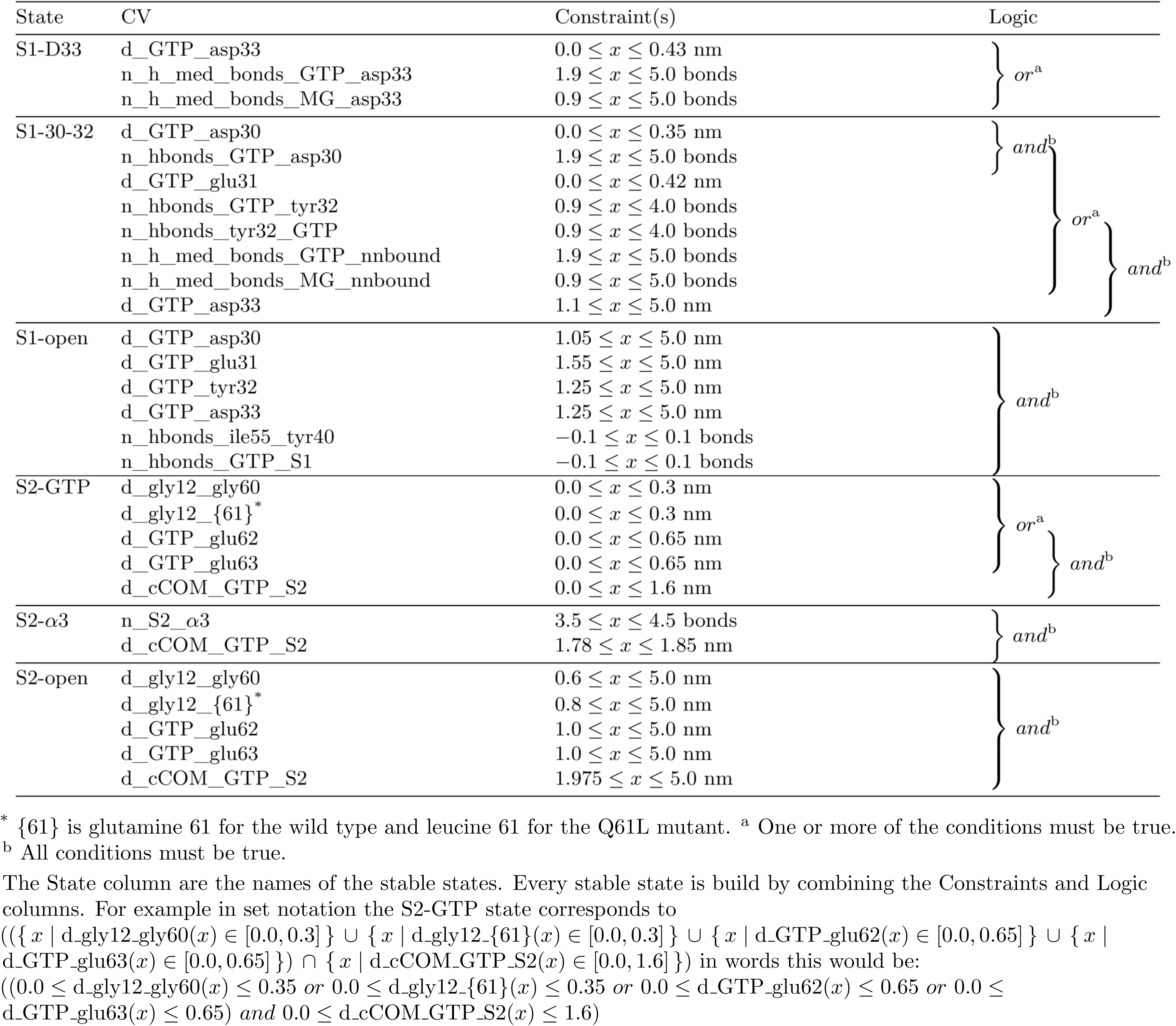
List of the stable state definitions for KRas.

#### Transition Path Sampling (TPS)

In the long molecular dynamics simulations some transitions spontaneously occurred. These transitions were used as the starting transition path for TPS [20, 37]. One TPS simulation was performed for S1, starting from the S1-30-32 to S1-open transition. For S2 three TPS simulations where performed, each starting from a different transition. This was done for both WT and Q61L. The initial trajectories were first equilibrated with a TPS simulation until the first decorrelated transition path (a transition path that has no frames in common with the original path) was obtained. This decorrelated path was used as the starting point for the production TPS simulations.

##### Settings

The TPS simulations were performed with OpenPathSampling(0.1.0.dev-c192493) [38, 39]. Multiple state TPS (MSTPS) [21] was performed with an all-to-all flexible length ensemble, excluding self-transitions. All-to-all means all transitions connecting two states are allowed. A self-transition is a path that starts in a states and returns to that same state after crossing the boundaries set by the state definitions. We used the one-way shooting algorithm [40], with uniform shooting point selection. For the S1 simulations, 1000 shooting trials were performed, while for each of the S2 simulations 2000 shooting trials were performed.

##### Analysis

All analysis of the TPS simulations was performed using the tools included in the OpenPathSampling package [38, 39], extended with custom Python code. Matplotlib [41] was used for plotting the graphs and triangles.

The supporting figures 8 and 9 show the type of transition as a function of the MC trial for S2 of WT and Q61L.

**Fig 2.**
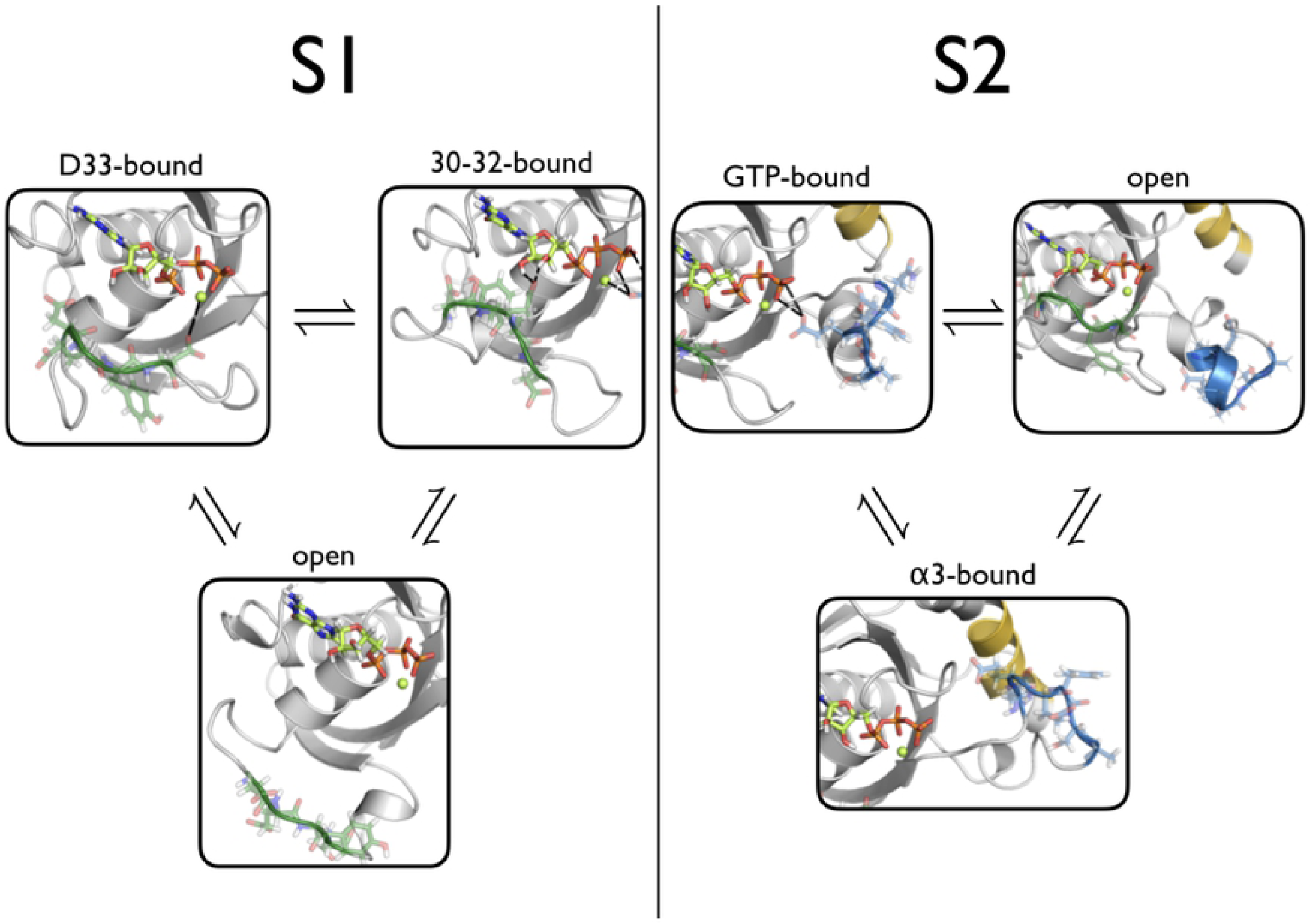
Stable states of KRas. The stable states found for S1 and S2 are shown with the same coloring as figure 1(left). The S1-D33 state corresponds to the conformation in which D33 in S1 has a hydrogen bond with GTP. The S1-30-32 state corresponds to the conformation in which one or more hydrogen bonds occur between residues 30-32 and GTP. The S1-open state corresponds to the conformation in which S1 has no interactions with GTP and is oriented away from GTP. For S2 the S2-GTP state corresponds to the conformation in which S2 has one or more hydrogen bonds with GTP. The S2-open state corresponds to a state in which S2 has no interactions with GTP and is oriented away from GTP. The S2-*α*3 state corresponds to a conformation in which S2 has no interactions with GTP, but instead has 4 interactions with the *α*3-helix.

**Fig 3.**
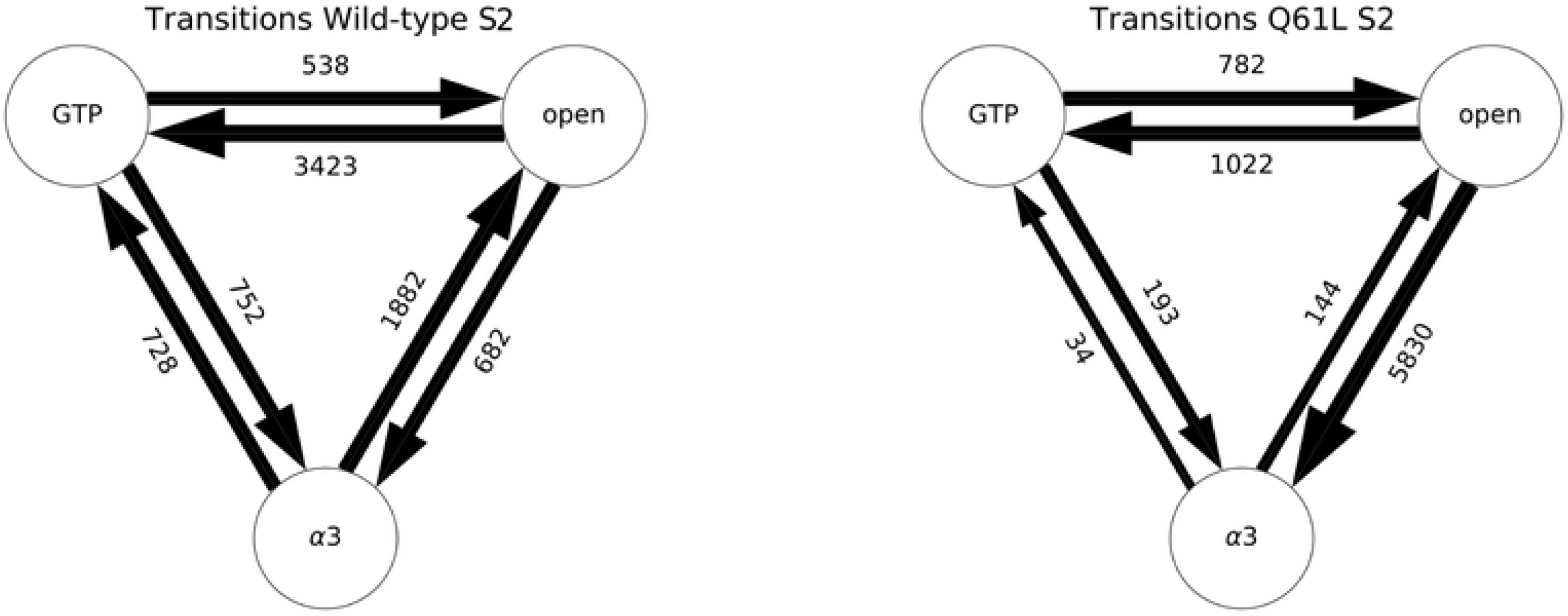
Transitions of S2 in WT and Q61L. A schematic representation of the number of samples per transition for the WT (left) and Q61L (right) TPS simulations. The circles correspond to the stable states of S2, with GTP being the S2-GTP state, open the S2-open state, and *α*3 the S2-*α*3 state. The arrows represent the sampled transition, pointing in the direction that was considered forward during the simulation. The labels are the number of accepted paths in each transition, and the width of the arrows is scaled with a 10 log scale of this number.

**Fig 4.**
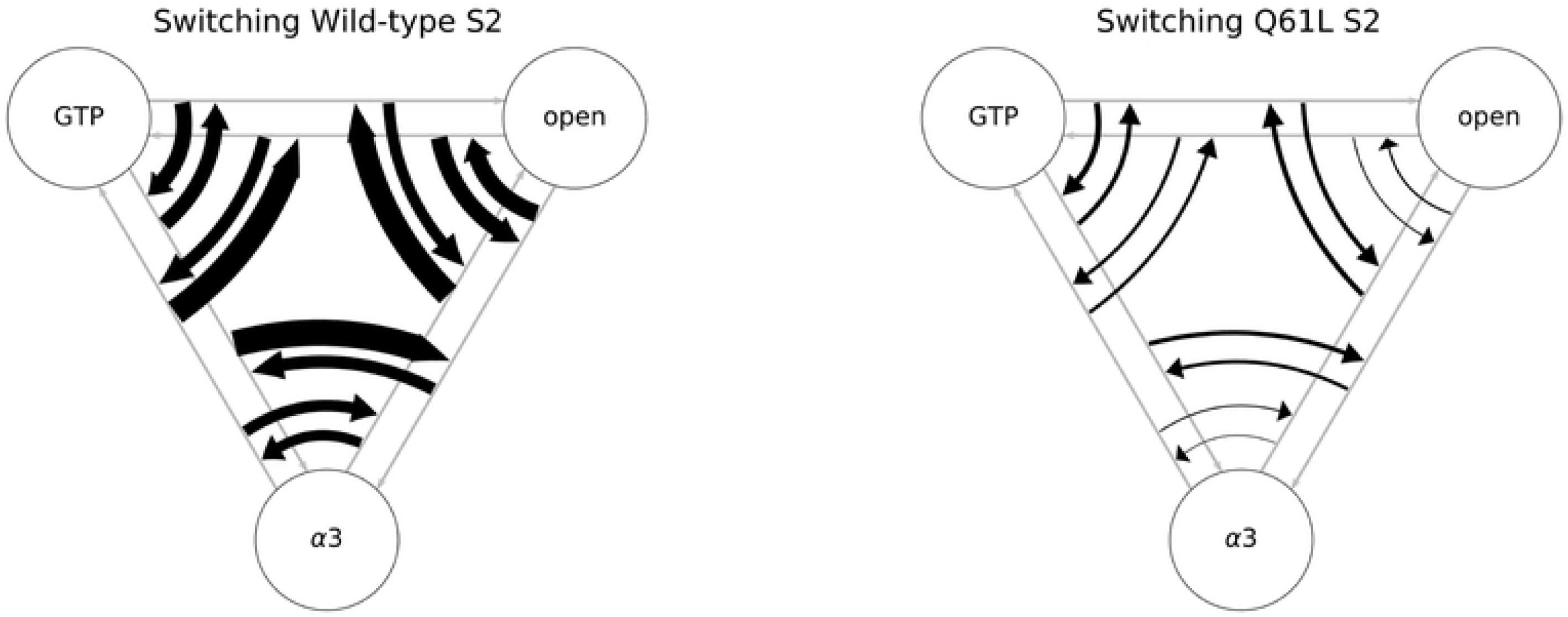
Switches between transitions. A schematic representation of the amount of switching between different sampled transitions in WT (left) and Q61L (right). The circles represent the stable states and the gray arrows show the unscaled transitions. The same state abbreviations were used as in figure 3. Each of the black arrows represents a switch between the transitions, scaled linearly to the number of times this switch occurred.

**Fig 5.**
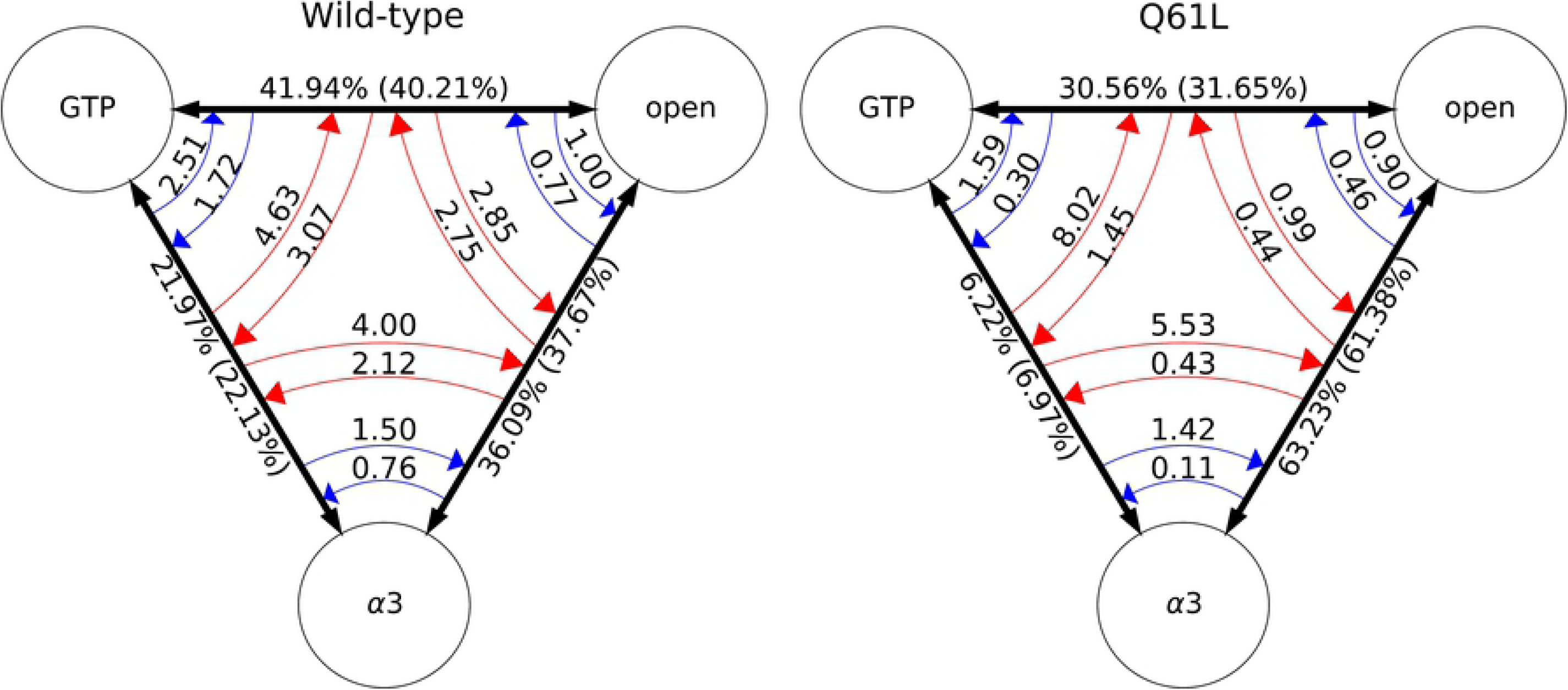
Population analysis. (left) All WT S2 simulations (right) Q61L simulations starting in the S2-*α*3 → S2-GTP transition and the S2-open → S2-*α*3 transition. The circles represent the stable states and the same state abbreviations are used as in figure 3. The black arrows are the time combined transitions, the red arrows are the switching rates obtained from all accepted paths, and the blue arrows are the switching rates observed from only using decorrelated paths. The labels of the transitions arrows are the population percentages from all accepted paths, with the decorrelated data in parentheses. The labels of the switching arrows are the rates of switching per 100 MC steps.

**Fig 6.**
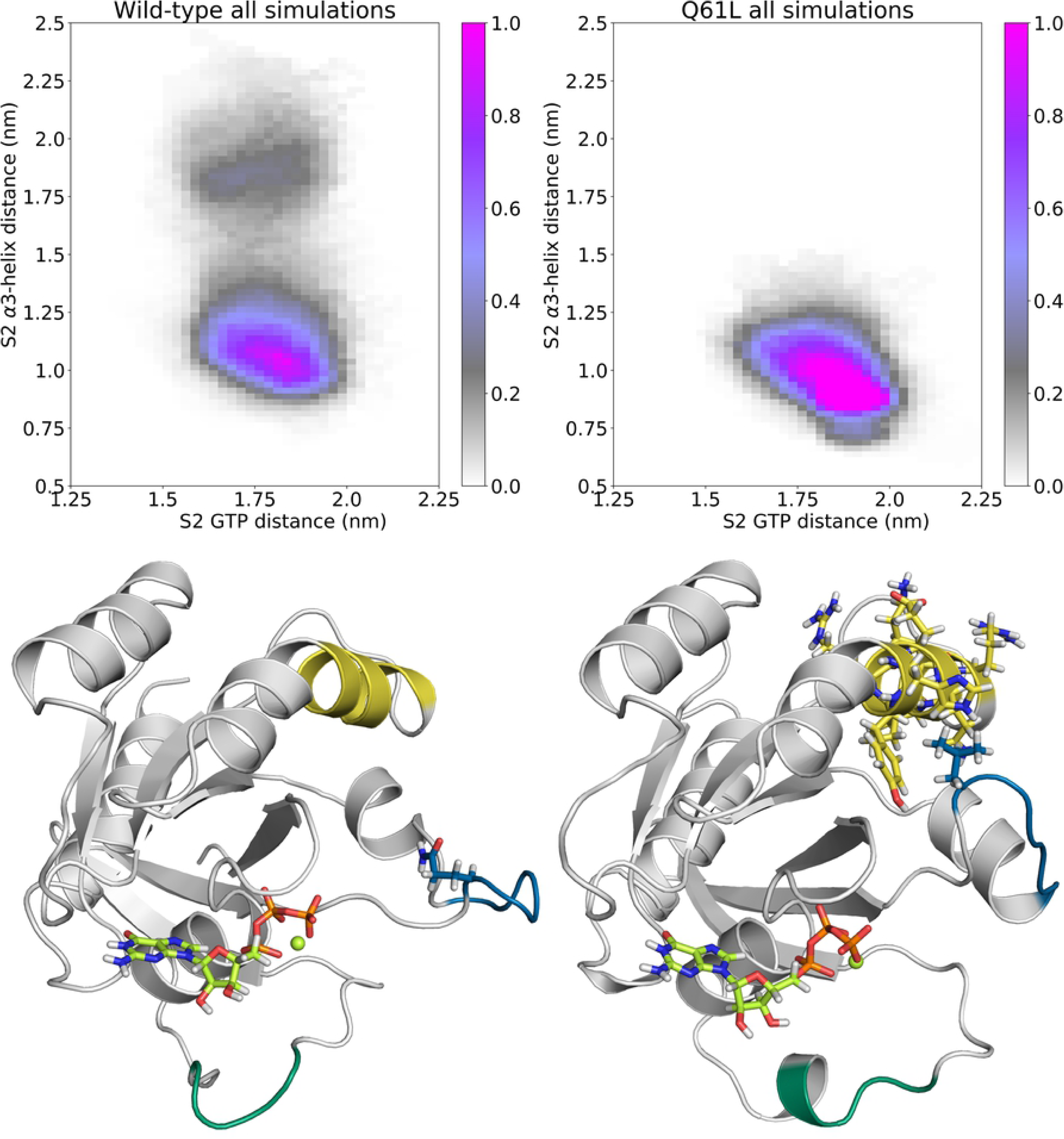
Mechanistic differences between WT and Q61L. (top) Path density histograms of the WT (left) and Q61L (right) simulations with on the x-axis the distance between the circular mean center of mass (cCOM) of S2 and on the y-axis the cCOM of GTP and on the y-axis the distance between the cCOM of S2 and the cCOM of residues 95, 96, 99, and 102. The bin widths are 0.25 Å for both axes. The coloring shows the sampled configuration of the transitions, weighted per trajectory, and normalized to 1. The stable states are not defined entirely based on these coordinates, but S2-GTP corresponds roughly to the area left of 1.6 nm on the x-axis, S2-open to the area right of 2 nm on the x-axis, and S2-*α*3 at intermediate distances on the x-axis, and below 0.8 on the y-axis. (bottom) Snapshots of the S2-open-state with the same coloring as figure 1(left) from (left) the channel that is far away from the *α*3-helix in the WT simulation and (right) the channel close to the *α*3-helix in the Q61L simulation.

**Fig 7.**
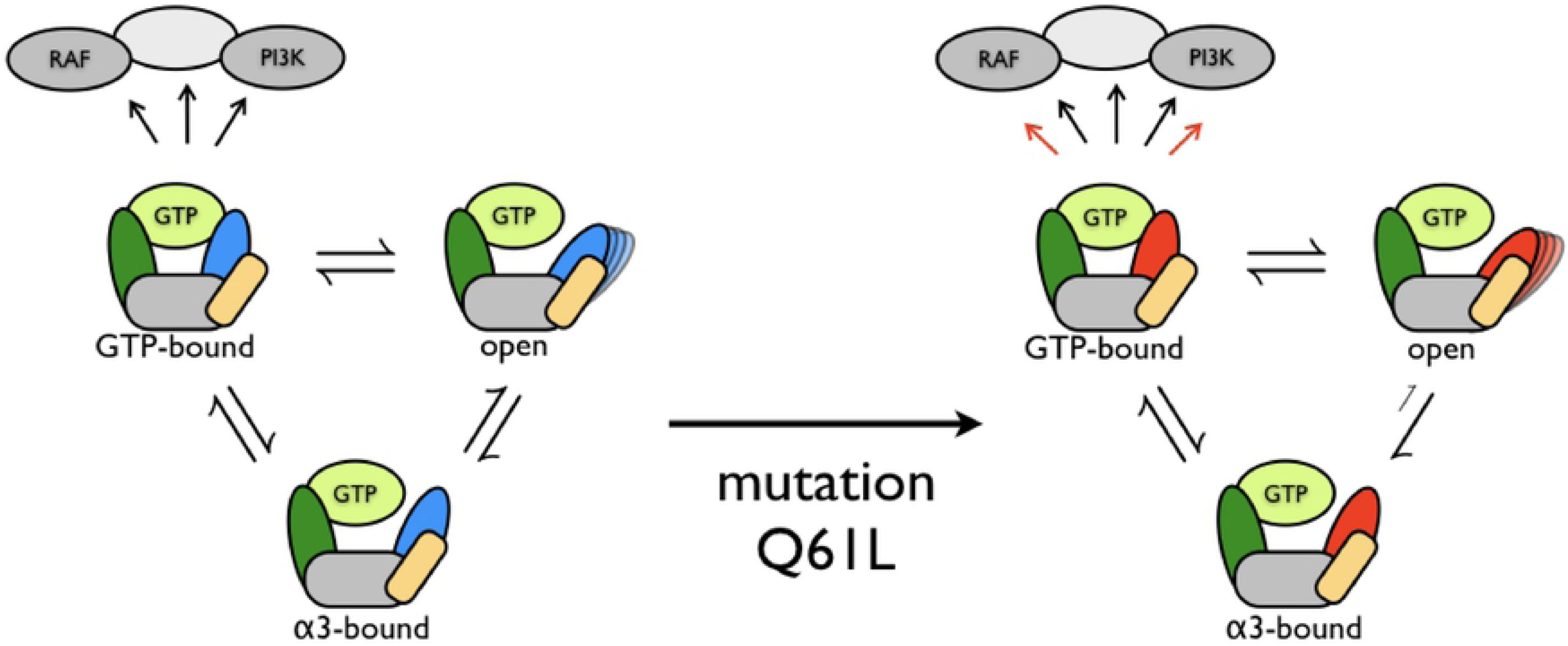
Schematic overview of the effect of the Q61L mutation on the dynamics of Kras. (left) WT (right) Q61L. The S1 region is represented in green, the S2 region in blue for WT and red for Q61L, the *α*3-helix in yellow, and the rest of the protein in grey. Downstream effectors are also shown in grey. Assuming only the S2 GTP-bound state triggers the downstream effectors, the Q61L mutation alters the conformational space such that one channel to reach the open state becomes very unlikely. This would lead to either a shift in the equilibrium distribution between the open and GTP-bound state or to transitions occurring more frequently. Both of these effects would lead to an increased probability to encounter downstream effectors while in the GTP-bound state, which would trigger the downstream signaling networks.

**Fig 8.**
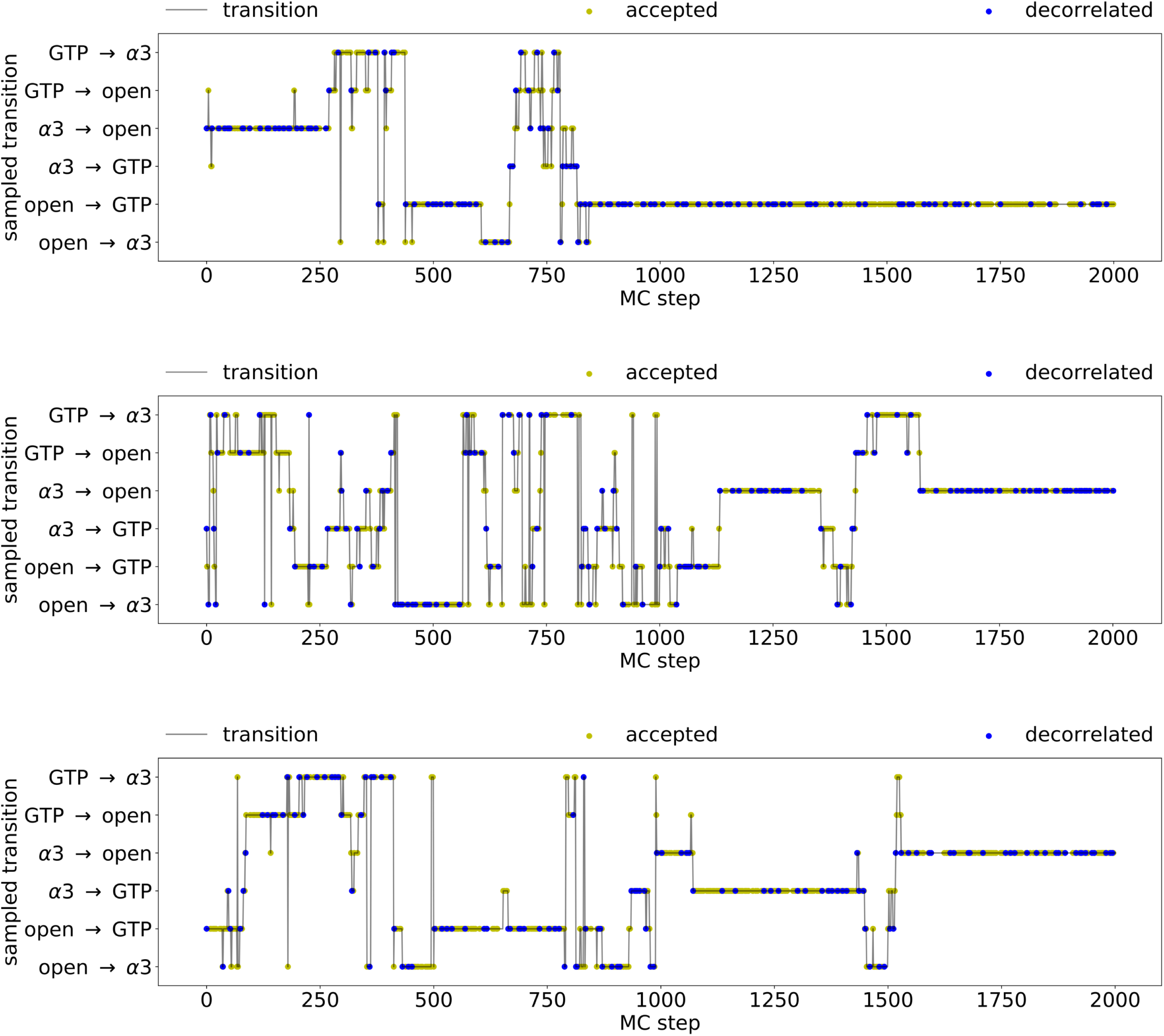
Transitions as function of the Monte-Carlo step for the WT simulations. The simulations started from (top) the S2-*α*3 to S2-open transition, (middle) the S2-*α*3 to S2-GTP transition and (bottom) the S2-open to S2-GTP transition. The x-axis shows the number of the MC steps. The y-axis shows the sampled transition. The y-axis lists all transitions that can occur for S2. The gray lines represents the trial moves, with the accepted MC steps highlighted as yellow dots and the accepted MC steps that lead to a new decorrelated trajectory with a blue dot.

**Fig 9.**
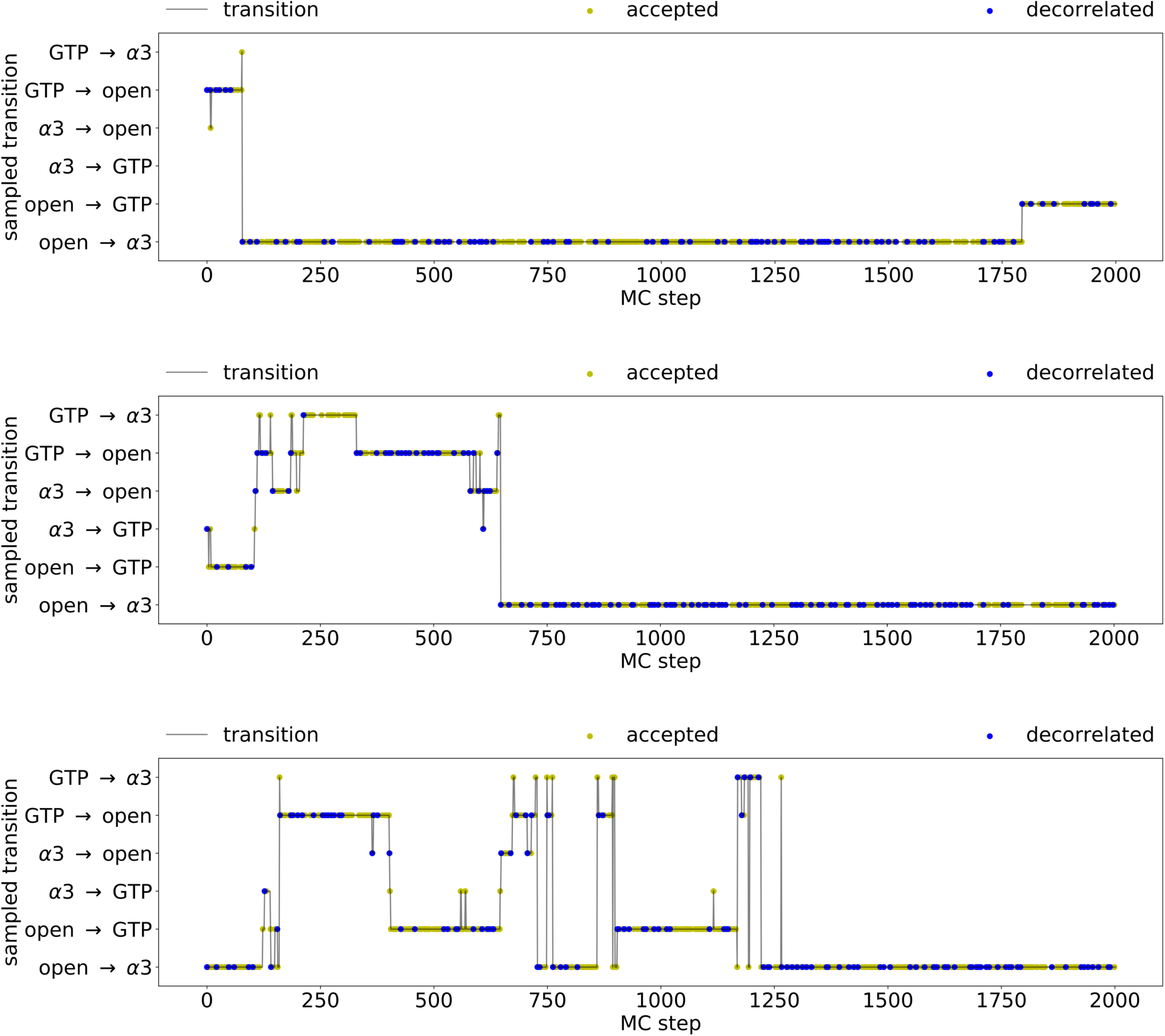
Transitions as function of the Monte-Carlo step for the Q61L simulations. The simulations started from (top) the S2-GTP to S2-open transition, (middle) the S2-*α*3 to S2-GTP transition and (bottom) the S2-open to S2-*α*3 transition. The x-axis shows the number of the MC steps. The y-axis shows the sampled transition. The y-axis lists all transitions that can occur for S2. The gray lines represents the trial moves, with the accepted MC steps highlighted as yellow dots and the accepted MC steps that lead to a new decorrelated trajectory with a blue dot.

##### Path density histograms

Path density histograms (pdhs) are two-dimensional histograms that show the configurations in a transition path, projected on collective variables. Each path is weighed with its MC weight, and divided by the number of total MC trials. For example, if a trajectory visits a histogram bin, the count of that bin is increased by the MC weight of that trajectory. It does not matter how often the trajectory visits a bin, it counts the trajectory only once.

##### Switching analysis

In MSTPS simulations, more than one transition is possible (e.g., *A* → *B, A* → *C, B* → *A*, etc.) However, only one simulation is sampled at a given MC step of the MSTPS simulation. The shooting algorithm allows the specific transition being sampled to change during the simulation. This helps provides heuristics for judging convergence. First, whether all transitions are visited give an estimate of the ergodicity. Second, forward and backward versions of transitions with the same pair of states (e.g., *A* → *B* and *B* → *A*) should have identical statistics in all ways. Furthermore, the fraction of MC steps spent in the two transitions between the same pair of states should be the same, as should the path length distributions.

With one-way shooting, switching between a transition *A* → *B* and its reversed version, *B* → *A*, requires at least three sequential switches: e.g., starting from an *A* → *B* transition, a transition from *A* → *C* can be generated, followed by a *B* → *C* transition, from which the next shot can result in a *B* → *A* transition. One-way shooting can only change the starting or ending state with a backward or forward shot respectively, but cannot change both in the same MC step. As MSTPS samples an equilibrium distribution, the number of paths collected from the *A* → *B* transition should be similar to the number of paths from *B* → *A*, reversed in time, which provides a measure for convergence of the simulation.

##### Kinetics analysis

As we assume that the switching samples an equilibrium distribution, the probability *P*_*i*_ of sampling a transition *i* is given by:

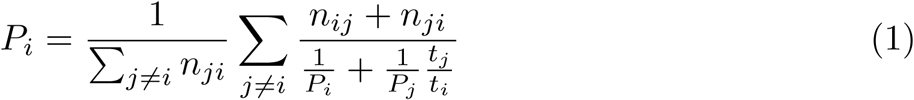

where *n*_*ij*_ is the number of switches from *i* to *j*, and *t*_*i*_ is the number of MC steps sampling transition *i*. As the sum of all probabilities is equal to one Σ_*i*_ *P*_*i*_ = 1, equation 1 can be solved for all *P*_*i*_. From the probabilities the switching rate from *i* to *j, k*_*ij*_, can be calculated by:

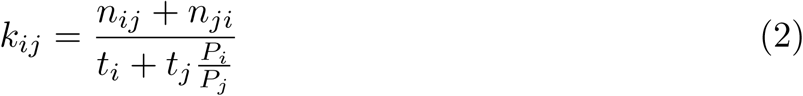

The values for *n* and *t* are taken from the MSTPS simulations. This analysis is adapted from [42].

## Results

### Identification of conformational states

The crystal structure of GppNHp bound HRas (PDB: 4EFL) [17, 18] was used as a structural template to model the sequence of WT and Q61L KRas-4B with GTP bound. With these two structures we performed four 100 ns MD simulations to explore the conformational space of KRas, for both WT and Q61L. These simulations resulted in the characterization of three stable states for S1, S1-D33, S1-30-32, and S1-open, and three stable states for S2, S2-GTP, S2-*α*3, and S2-open, shown in figure 2. When S1 is in the S1-D33 state, the side chain of D33 is involved in (water-mediated) hydrogen bond interactions with GTP. For the S1-30-32 state, S1 has shifted along GTP, compared to the S1-D33 state, to form one or more hydrogen bonds between the side chains of residues D30, E31, or Y32 and GTP. The conformations in which S1 has no hydrogen bond interaction with GTP and where it is oriented away from GTP are classified as the S1-open state. The S2-GTP state corresponds to the conformation of S2 where it forms hydrogen bonds with GTP. Two states can occur when S2 is oriented away from GTP. In the S2-*α*3 state, S2 has multiple interactions between its side chains and the *α*3-helix. In the S2-open state S2 has no binding interactions with GTP. The parameters for defining these states are listed in Appendix 1. Within the timescale of the path sampling simulations, the conformation of S1 has little effect on the conformation of S2 or vice versa. The same stable states were found for both WT and Q61L, and were stable for at least 100 ns of MD.

### Mapping conformational transitions

Using MSTPS, we investigated the transitions between the stable states as identified in the MD simulations for S1 and S2 separately. For both S1 and S2 three pairs of transitions can occur: S1-D33 ↔ S1-30-32, S1-D33 ↔ S1-open, and S1-30-32 ↔ S1-open for S1, and S2-GTP ↔ S2-*α*3, S2-GTP ↔ S2-open, and S2-*α*3 ↔ S2-open for S2. For both WT and Q61L we performed one MSTPS simulation for S1 starting at the S1-30-32 ↔ S1-open transition, and three independent MSTPS simulations for S2, each starting in a different transition, resulting in eight MSTPS simulations in total. The statistics of the MSTPS simulations are listed in table1 and indicate a good acceptance ratio of 34% or higher, and an aggregate simulation time of microseconds.

Path sampling simulations for proteins (with stochastic dynamics and diffuse barriers) commonly employ the stochastic, or “one-way” shooting algorithm [37], which improves the acceptance ratio. In this algorithm a trial move replaces only part of the trajectory (forward or backward). Therefore, successive trajectories will have segments with overlapping frames and at least two trials (one forward and one backward) are needed for an accepted trajectory to have no frames in common with the original. These no-overlap trajectories are referred to as “decorrelated”, and are required for sufficient sampling. Although MSTPS only samples one transition at a time, switching between transitions can occur when there are more than two states. For example, given states *A, B*, and *C*, an initial *A* → *B* trajectory can produce a trial *A* → *C* trajectory if a forward trial ends in state *C*. Such a transition of transitions is called a “switch.” Analyzing the switching behavior provides useful insight into the transition region.

The transitions sampled in the MSTPS simulations of S2 are summarized in figure 3. Between each pair of states there are two possible transitions, corresponding to what is the forward time direction in the path. Both the WT and the Q61L simulations have sampled all allowed transitions. At infinite sampling, the relative sampling frequency for the two transitions between a given pair of states will be identical. While different, the counts for most transition pairs are within an order of magnitude of each other, indicating acceptable sampling. The exception is the S2-open and S2-*α*3 pair in Q61L, where the S2-open→S2-*α*3 transition is sampled 5830 times, and the S2-open←S2-*α*3 transition only 114 times. This difference in sampling the S2-open and S2-*α*3 transition in WT and Q61L suggests that the mutation has altered the transition region.

Analyzing the switching behavior can provide insight into the transition region. A lack of switching between two states indicates there is a large (free energy) barrier in the transition region between the channels for the individual reactions. Conversely, many switching events suggests a flatter, more diffusive landscape in the transition region. The figures in appendix S2 plot the transition sampled in each accepted path as a function of the number of MC steps for all S2 simulations. For the WT simulations MC steps resulting in accepted trajectories and in decorrelated (no-overlap) trajectories are distributed uniformly throughout all 3 simulations, indicating that none of the simulations was trapped in a part of conformational space from where the probability of accepting a new (part of a) path is exceedingly low. Throughout most of the WT simulations, switching occurs on average every 16 MC steps, indicating that the simulation loses memory of the starting transition path. The second part of the simulation starting from an S2-*α*3 to S2-open transition is an exception, as this simulation remains in the S2-open → S2-GTP transition for over 1000 MC steps. For the Q61L simulations the accepted and decorrelating MC steps are also distributed uniformly throughout all three simulations. All simulations spend a significant amount of simulation steps in the S2-open → S2-*α*3 transition, possibly indicating that the open state of Q61L is close to the S2-*α*3 state. The number of switches occurring between the transitions is much lower than in the WT simulations. The lack of switching also explains why the Q61L simulations are less well sampled than the WT simulations. Figure 4 shows the number of switches between the six transitions occurring for S2 as arrows with a thickness relative to the number of switches. Clearly, the number of switches between transitions is much lower for the mutant than for the WT, indicating that the WT has a lower free energy barrier between the different transition channels than Q61L.

### Kinetic analysis of switching

To quantify the relative frequency or “population” of each transition and the switching rate between transitions, we applied a kinetics analysis approach as developed for Replica Exchange MD [42] on our MSTPS data. In this analysis, rate constants are estimated from the number of transitions between stable states and the average residence times in the stable states. To apply this analysis to our MSTPS data, time, stable states and transitions correspond to respectively the MC trials, the transitions between the states and the switches (transitions of transitions). Such a kinetics analysis results in rate constants that measure the switching rate between transitions in units of MC steps, see the Methods and Materials section for more detail. We analyzed the kinetics by including all accepted trajectories, or only decorrelated trajectories, see Figure 5. For one Q61L simulation, starting from S2-GTP → S2-open, no switching occurs out of the S2-GTP ↔ S2-*α*3 transition and we therefore excluded this simulation from the switching analysis. The “population” of the S2-*α*3 ↔ S2-open transition is almost twice as high for Q61L (63.23%) than for WT (36.09%), while the S2-GTP ↔ S2-*α*3 is less likely for Q61L (6.22%) than for WT (21.97%). When including all accepted paths, most switching rates are lower for Q61L than for WT, except for the switches out of the S2-GTP ↔ S2-*α*3 transition.

The largest relative difference between WT and Q61L are observed for the switching rates out of the S2-*α*3 ↔ S2-open transition. When including only decorrelated trajectories, all switching rates are lower for Q61L, with the largest relative difference for the transition into the S2-GTP ↔ S2-*α*3 transition. These observations indicate that the S2-*α*3 and S2-open states are closer together in Q61L than in WT. In addition, it is difficult to obtain decorrelated paths for the S2-GTP ↔ S2-*α*3 transition. This is also apparent from the figures in Appendix S2, where the residence time in the S2-GTP ↔ S2-*α*3 transitions is almost always only a single MC step. Less switching would occur if the transition channels are narrower and thus have a steeper slope in free energy orthogonal to the reaction channel. Alternatively, part of the region between transitions is less favorable for Q61L compared to WT.

### Path densities reveal two channels

To further investigate the origin of the difference in switching kinetics, we projected the trajectory space sampled in the combined TPS simulations in a path density histogram (see the Methods section for an explanation on how path density histograms are computed). Figure 6(top) shows the path density for the WT (left) and Q61L (right) S2 TPS simulations, projected in the plane of the distances between S2 and GTP, and between S2 and residues H95, Y96, Q99, and R102 of the *α*3-helix (see Appendix S1for definitions of these distances). Note that path densities do not show stable states, as the trajectories are stopped when reaching one. Comparing the two path density plots indicate that the WT simulations sample a larger region on the y-axis, as the WT path density extends to above 1.25 nm, while the Q61L path density is more confined to the region below 1.25 nm in *S*2 − *α*3 distance.

Inspection of the least changed path, a trajectory connecting paths on top of the transition barrier, in figures 10 and 11 in Appendix S3, confirms that the observed switching is indeed a diffusive process and that the Q61L mutation constrains the dynamics. This indicates that S2 can move away from the *α*3-helix more easily in the WT protein. The WT histogram even shows a second channel for transitions between the S2-GTP and the S2-open states, at a distance of more than 1.75 nm from the *α*3-helix. The Q61L simulations sample configurations closer to the *α*3-helix, as indicated by the density below 0.8 nm on the y-axis. A more pronounced negative correlation exists between the S2 - *α*3-helix and the S2 - GTP distances. The further away S2 is from GTP, the closer it is to *α*3. Furthermore, the Q61L simulations do not sample the second channel at all. As three independent simulations were performed for both WT and Q61L, each initiated from a different transition, the absence of direct solvation transitions for Q61L are likely to be a direct consequence of the mutation.

**Fig 10.**
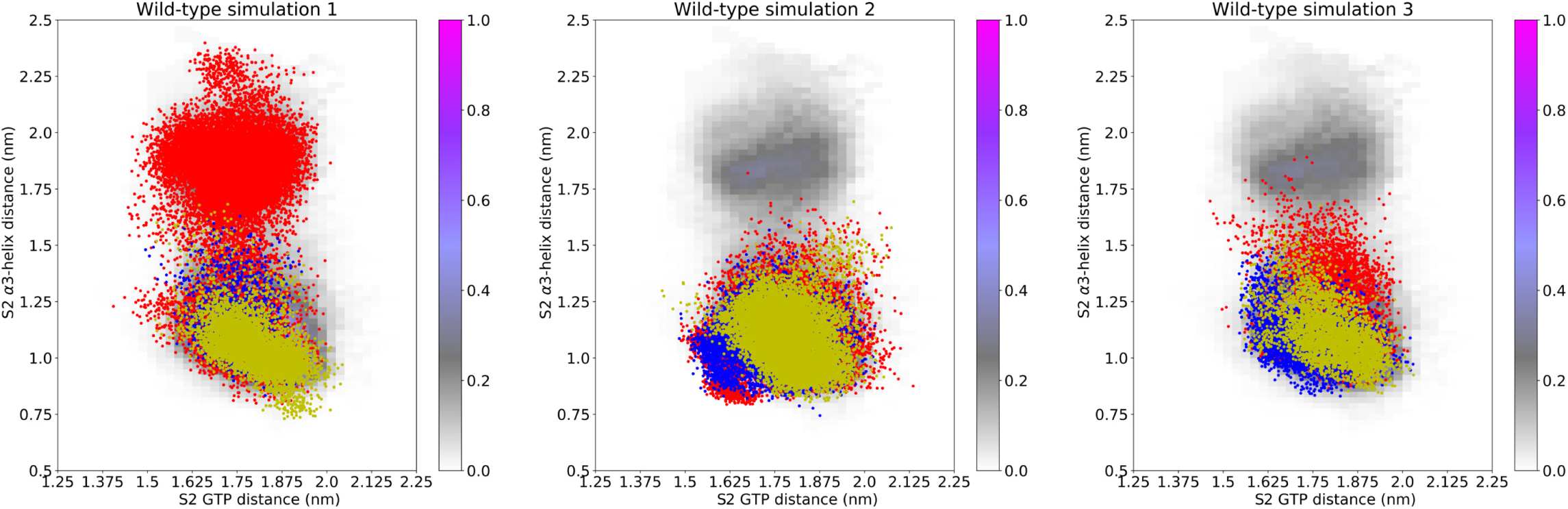
The Least-Changed-Paths of the WT simulations. The frames of the LCP of each WT simulation, shown on top of the combined path density histogram, as shown in (figure 6(top)). The color of each frame represents the first transition sampled by that frame, red for S2-GTP ↔ S2-open, blue for S2-GTP ↔ S2-*α*3, and yellow for S2-*α*3 ↔ S2-open. The numbering of the simulations is in the order of figure 8.

**Fig 11.**
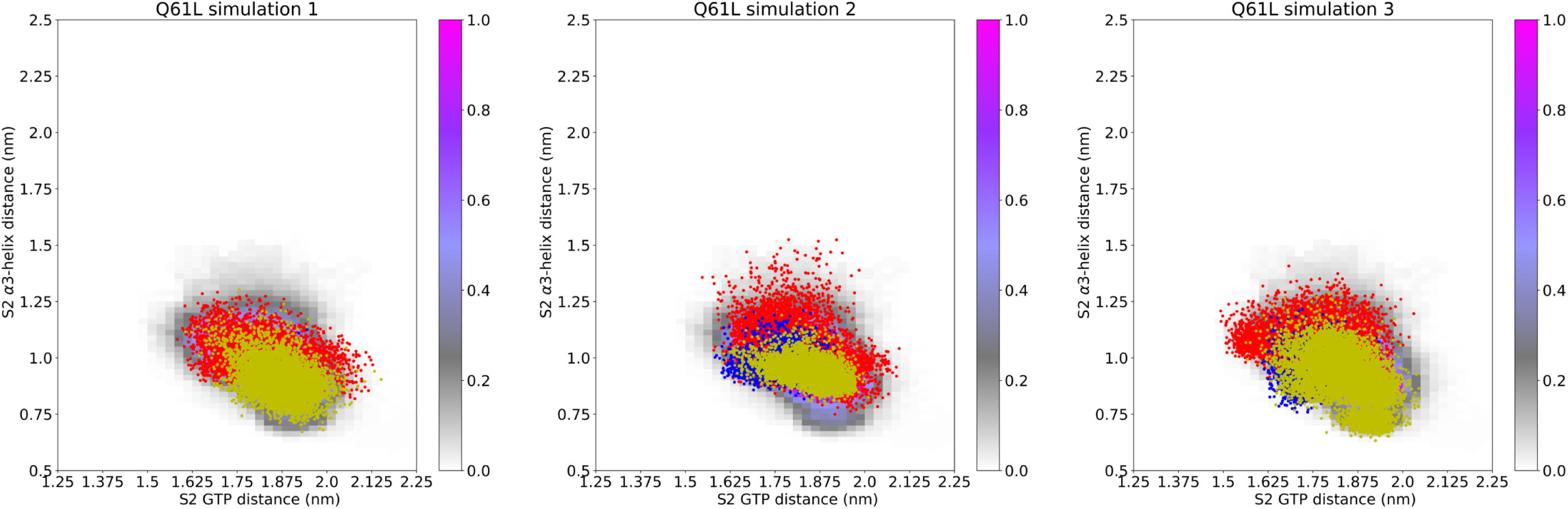
The Least-Changed-Paths of the Q61L simulations. The LCP are shown on top of the combined path density histogram (figure 6(top)). The color of each frame represents the first transition sampled by that frame, and is the same as in figure 10. The numbering of the simulations is in the order of figure 9.

The two conformations plotted in figure 6(bottom) illustrate the difference between the two reaction mechanisms or channels. The image on the left shows the WT protein in the S2-open state with S2 at a distance of at least 1.75 nm from the *α*3-helix. On the right Q61L is shown in the S2-open state with S2 closer than 1.25 nm to the *α*3-helix, see video supplements 1 and 2for movies of typical trajectories for each of the two reaction channels. The distance of S2 to *α*3 is indicative of the different mechanisms. The channel far away from the *α*3-helix represents a mechanism involving water molecules solvating S2, resulting in S2 extending into the solvent, away from both GTP and the *α*3-helix. The channel close to the *α*3-helix represents S2 moving along a hydrophobic pocket on the *α*3-helix. In this reaction mechanism, S2 can either enter the S2-*α*3 state by forming four contacts between S2 and the *α*3-helix, or by sliding along the helix until entering S2-open. The Q61L mutation changes a hydrophilic residue to a hydrophobic one, thus lowering the affinity of S2 for water. Therefore, this latter, solvated, channel, which is easily accessible for the WT protein, becomes much less likely for Q61L. Moreover, the mutated S2 has stronger interactions with the *α*3-helix, as shown by the higher path density in the channel close to *α*3-helix (Figure 6(top, right)), indicating that for Q61L it is harder to escape from the *α*3-state. The increased stability of the *α*3-state renders the S2-GTP↔S2-open transition less likely. Figure 7 summarizes this conclusion.

### Q61L has a higher propensity for a more structured S2-open

Visual inspection of the transition paths shows that in some WT trajectories the *α*2-helix (residues 65–73, overlapping with part of S2), unfolds when entering the S2-open state, but retains its shape for the Q61L mutant. Probability histograms of the S2-open state obtained from the transition path ensemble by projection on the number of helical hydrogen bonds in the *α*2-helix and the S2-*α*3-helix distance shown in Appendix S3Figure 12 further substantiates this observation. Therefore, we can conclude that the S2-open state contains multiple sub-states, characterized by the conformation of the *α*2-helix and the S2-*α*3 distance. Furthermore, these probability histograms show that Q61L has a higher propensity compared to WT for the more structured conformations of the S2-open state. The *α*2 helix plays a vital role in binding other proteins [7], suggesting that these structured sub-states are more similar to the active state 2 than the inactive state 1.

**Fig 12.**
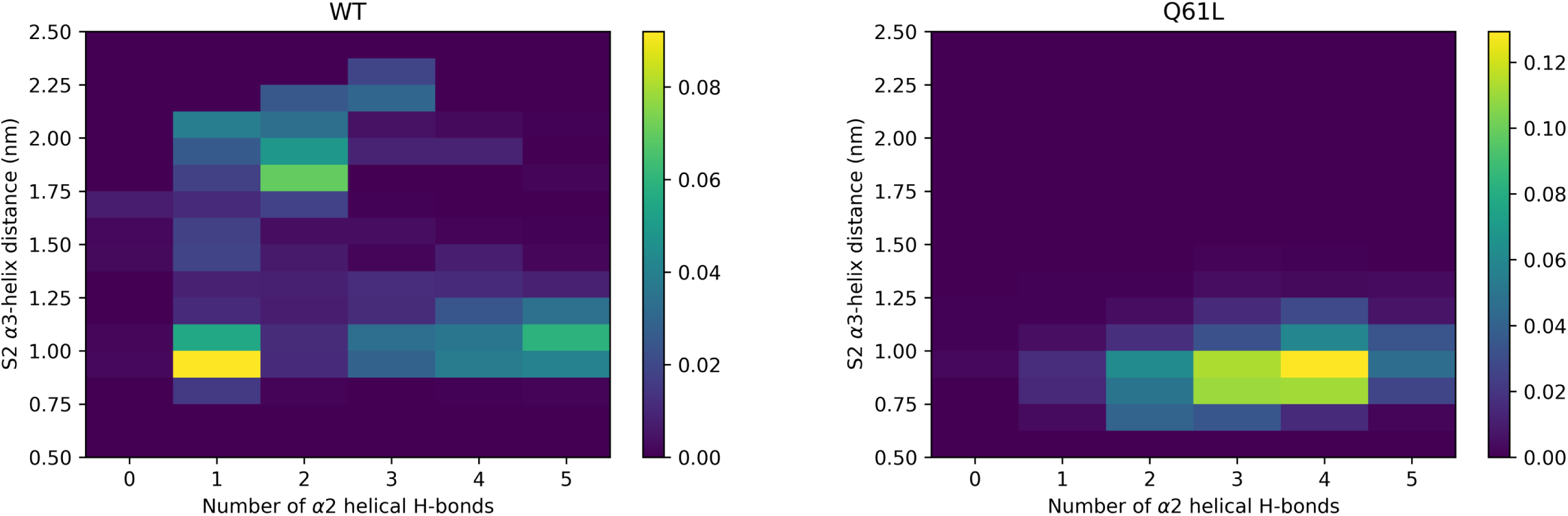
Two-dimensional probability histogram of the S2-*α*3-helix distance and the number of helical hydrogen bonds in *α*2-helix for the S2-open state. These are shown for (left) WT and (right) Q61L. The y-axis is the cCOM of S2 to the *α*3-helix (as used in figure 6). The x-axis is the number of hydrogen bonds (as described in table 2) between the backbone O of residue *i* and the backbone NH of residue *i* + 4 for *i ∈* [65, 69]. The colors indicate the probability.

**Fig 13.**
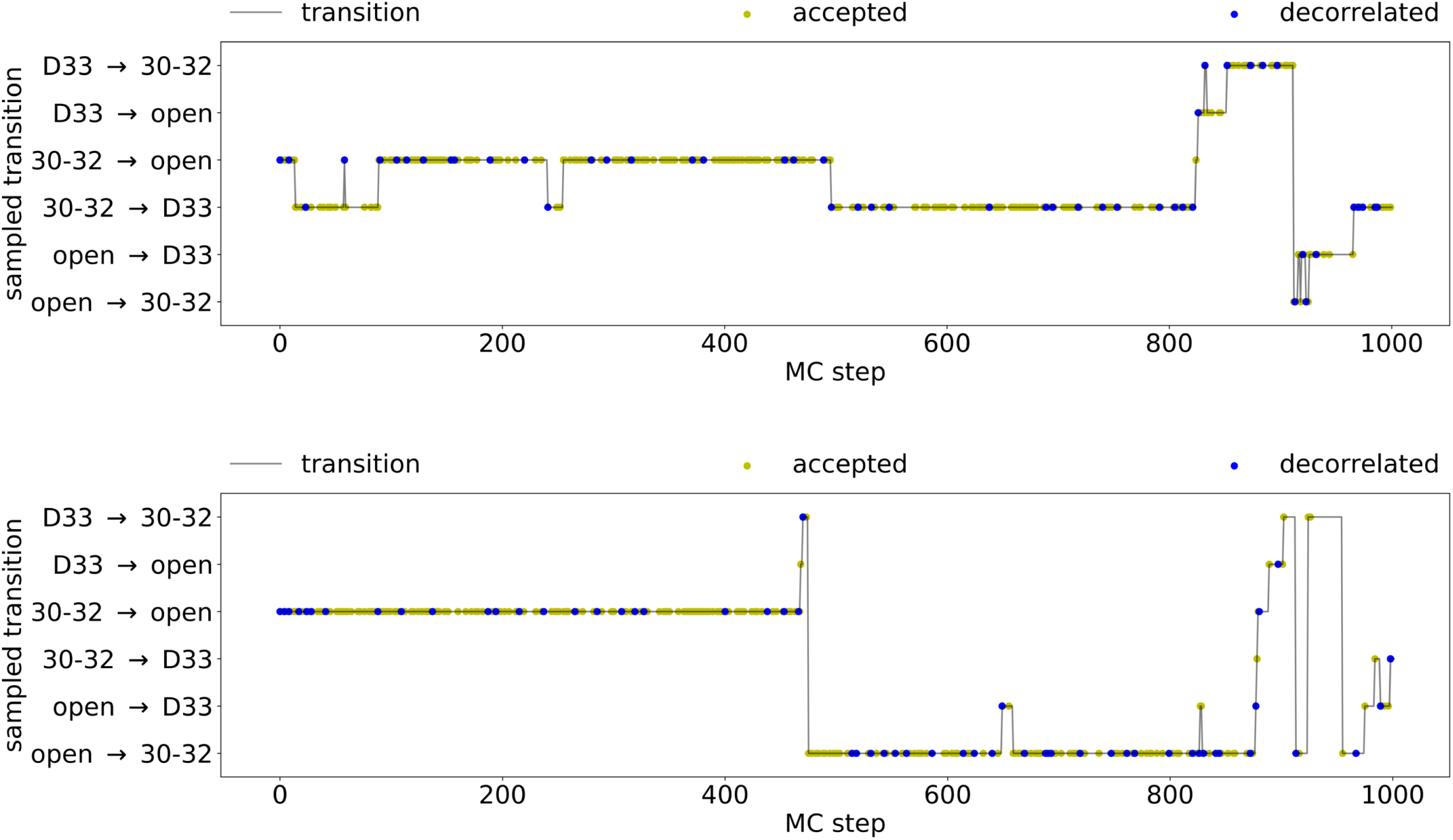
Transitions as function of the Monte-Carlo step for the S1 simulations. (top) WT (bottom) Q61L. The same axis setup and labeling is used as in the supplements for figures 8 and 9. Here D33 corresponds to the S1-D33 state, 30-32 to the S1-30-32 state, and open to the S1-open state.

**Fig 14.**
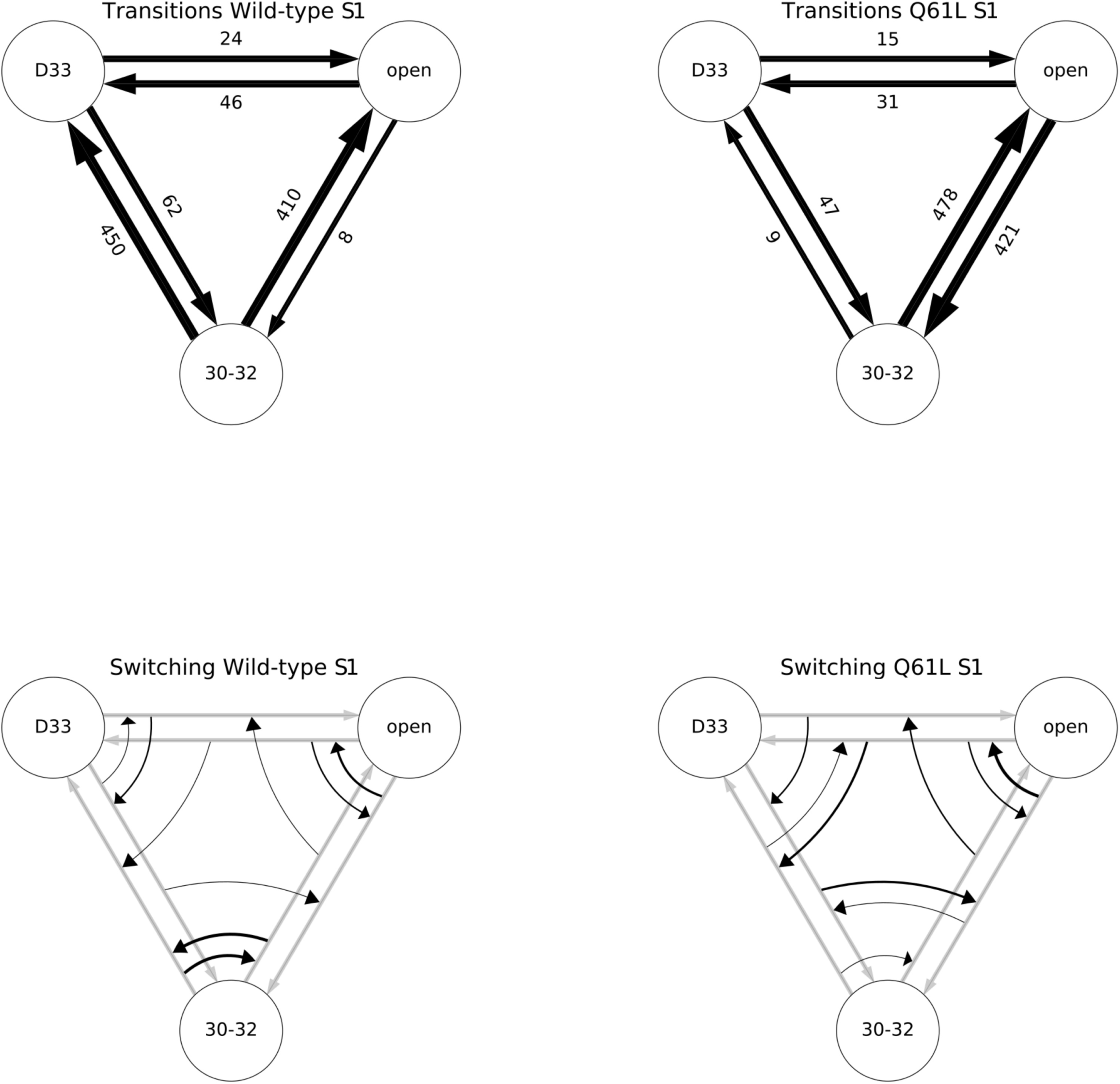
Results of the S1 path sampling simulations. (left) Transitions in the (top, left) WT and (bottom, left) Q61L simulations. (right) Switches in the (top, right) WT and (bottom, right) Q61L simulations. The same labeling for the states is used as in figure 13.

The more open and flexible sub-states of the S2-open state are less likely to be recognized by downstream effectors. A *β*-strand in the PI3 kinase interacts with KRas via both S1 and S2 [6], which can only occur when both S1 and S2 are in a closed conformation. The more open conformations are harder to reach in Q61L, and indeed, we only observe these flexible sub-states in the WT simulations, thus providing an explanation for the increased probability of Q61L to bind a downstream effector. This is consistent with the fact that the Q61L mutation is more prevalent in tumors involving HRas than KRas. Residue 95 is a glutamine in KRas, whereas it is a histidine in HRas, rendering the *α*3 helix slightly less hydrophilic and thus potentially enhancing the interaction with the mutated S2. This prediction may be tested by repeating the NMR experiment as performed by Geyer et al. [11], comparing the effect of the Q61L mutation on the switching frequency. Alternatively, the lifetime of the S2-*α*3 state could be measured using ^15^N NMR spectroscopy, by labeling nitrogen atom NE2 in the Q95 side chain, located in the *α*3-helix. Finally, the *α*3-helix identified as important for the transitions between the ordered, active state 2 and the flexible, inactive state 1 might provide a new target for the development of compounds that could ameliorate the effect induced by the Q61L mutation.

## Conclusion

In this work we investigated the conformational space and dynamic behavior of KRas in complex with GTP using path sampling. The loops in KRas that interact with GTP each visit three different conformational states. Surprisingly, these conformational states do not change upon introducing the Q61L mutation, located in region S2. However, the mutation has a significant effect on the transitions between the conformational states of region S2. While the WT protein frequently changes from one transition to another, the mutant hardly changes at all. Closer examination of the various transitions revealed that S2 in the WT protein is more likely to be solvated than in the Q61L mutant. The Q61L mutation prevents direct solvation of S2, which is an accessible route for the WT protein. Both WT and mutant can reach the opened up state by S2 sliding along a slightly hydrophobic pocket of the *α*3-helix. Reaching the open state may involve unfolding of the *α*2-helix, which hardly occurs in the Q61L system. Our results show that the TPS methodology in combination with the novel switching analysis, is able to map out the dynamics of a Ras protein, indicate differences in dynamics between the WT protein and an oncogenic mutant, and reveal details on the nature of the altered behavior as caused by the mutation. As such, our approach presents a powerful tool in providing a better understanding of the dynamic nature of Ras proteins, and may potentially aid in identifying new therapeutic targets for tumors induced by Ras mutations.

## Supporting information

**Appendix S1: Stable state definitions** The stable states are defined by ranges in collective variables. This appendix provides a guide to these stable state definitions. Table 2 gives the types of collective variables, while Tables 3 and 4 list the collective variables used to define the stable states for S1 and S2, respectively. Table 5 gives the ranges in collective variable space for the stable states found for S1 and S2.

**Appendix S2: Transitions as function of the Monte-Carlo (MC) steps** The figures listed in this appendix show the type of transition as sampled for each step in the TPS simulations.

**Appendix S3: MSTPS results for S2** The sampling statistics of the S2 MSTPS simulations are shown in table 1. The number of Monte Carlo (MC) trials was equal for all simulations. The acceptance is between 34 % and 42 %, which is reasonable considering the theoretical maximum of 67 %. This theoretical maximum is due to the fact that in our shooting algorithm only self transitions are forbidden. This leads to a maximum acceptance of 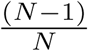, for *N* number of states. With *N* = 3 this leads to the theoretical maximum acceptance of 67% for this MSTPS study. The number of decorrelated trajectories is satisfactory for all simulations, and are spread well throughout the simulation as shown by the blue dots in the figures in Appendix 1. The average path length and total simulation time are only different for WT simulation 1. This simulation enters a different transition channel than the other simulations, which would explain these altered numbers.

The Least Changed Path (LCP) is the sequence of frames that, together, represent all accepted trajectories of a path sampling simulation. When running backwards in the simulation (from the last MC step to the first) a sequence of frames that are in between the shooting point of the latest trajectory and the shooting point of the trajectory in which that shooting point of the latest trajectory is replaced, on the last accepted trajectory before this next trajectory is added to the LCP. This is continued until the first MC step is reached. These LCPs represent the barrier region that is sampled during the TPS simulation. [45]. Figure 10 (WT) and figure 11 (Q61L) show the LCPs for all simulations, projected on top of the combined pdhs from figure 6. The colouring is based on the first sampled transition of each frame of the LCP and is red for S2-GTP S2-open, blue for S2-GTP ↔ S2-*α*3, and yellow for S2-*α*3 ↔ S2-open. For the WT, all transitions sample the same diffuse barrier region, as indicated by the overlap of the clouds, which supports the hypothesis that the switching is also a diffusive process. For the extra channel for the S2-GTP↔S2-open transition, this mostly occurs in simulation 1 of WT, but it is also observed in simulation 2 and 3. For the Q61L simulations, the LCP is more constrained to a value of under 1.3 nm for the S2-*α*3-distance. Also, the clouds overlap less with each other, making switching more unlikely. Simulation 1 and 3 of Q61L also show sampling of an extra S2-*α*3 ↔ S2-open channel, at values of 0.75 nm or lower for the S2-*α*3-distance.

Visual inspection of the transition paths as sampled for WT shows that in some paths helix *α*2 (residues 65-73) contained within S2, unfolds when entering the open state, but retains its shape in the S2-open state for the Q61L mutant. Two-dimensional probability histograms of the S2-*α*3 distance and the number of helical hydrogen bonds of the *α*2-helix (residues 65-73), for frames in the S2-open state, are shown in figure 12 for both WT and Q61L. Looking at the WT plot, there are two maxima for states in the reaction channel close to the *α*3-helix (under 1.5 nm on the y-axis), one where the *α*2-helix has all 5 helical H-bonds and one where it has only 1 helical H-bond. For the WT S2-open states away from the *α*3-helix (above 1.5 nm on the y-axis), the *α*2-helix has lost part of it helical structure, as indicated by a distribution around 2 helical H-bonds. When looking at the transition region between these two reaction channels at around 1.5 on the y-axis, helix *α*2 has lost most of its helical hydrogen bonds, which may indicate a correlation between the unfolding of helix *α*2 and the switching between the two reaction channels. The probability histogram for the S2-open frames of Q61L show a maximum at 4 helical H-bonds and S2 close to helix *α*3. These observations suggest that Q61L has a more structured open state.

**Appendix S4: MSTPS results for S1** The transitions as function of the MC trials of the S1 simulations are shown in figure 13. The x-axes represent the number of the MC trials, while the y-axis shows the sampled transition. Like in the supplements of figure 3, the y-axis lists the transitions, ordered such that the simulation can only switch to the transitions directly above or below the current transition, or between the top and bottom transition. This ordering is possible, because only either the initial or the final state can change per MC step due to the one-way shooting algorithm. With a forward shot the final state can change and a backward shot may change the initial state. The trial moves are shown as a gray line, with the accepted MC steps highlighted as yellow dots and the accepted MC steps that result in a new decorrelated trajectory with a blue dot. The accepted and decorrelating MC steps are distributed well throughout both simulations. The number of switches that occur between the transitions is similar for both WT and Q61L. Both simulations spend a significant amount of simulation steps in the 30-32 → open transition.

The sampling and switching behaviour of both simulations is summarized in figure 14. When looking at the transitions sampled for WT, the number of samples differ for the directions of the 30-32 ↔ D33 and the 30-32 ↔ open by more than an order of magnitude, this indicates that this simulation did not converge. For the Q61L S1 system, the number of samples in both directions of each transition is within the same order of magnitude, which is a first indication that this simulation might be converged. No large differences occur in the sampling during the WT and the Q61L simulations, as for all three transitions the WT has at least one direction with the same order of magnitude as the Q61L (like the D33→30-32 transition, which has 62 an 47 samples for respectively the WT and Q61L). However, for the open→30-32 and 30-32→D33 transitions the number of samples differ, with 8 for WT and 421 for Q61L and 450 for WT and 9 for Q61L.

This could be attributed to a lack of convergence of the simulations. The number of switches that occur between the transitions is similar for both the WT and the Q61L simulation.

Representative trajectories of all sampled transitions of both the WT and the Q61L were visually compared. No distinct differences in transition mechanisms between WT and the Q61L were observed. In conclusion, these results suggest that the mutation in S2 has little effect on the dynamical behavior of S1.

**S5 Video Movie of a typical trajectory of the S2-GTP and the S2-open transition for WT.** The frames in these movies are rendered with the switch regions highlighted in green for S1 and blue for S2. The *α*3-helix is highlighted in yellow. The protein is shown as a ribbon with an transparent stick representation for the amino acids in S1 and S2. GTP is shown as solid sticks, with carbon atoms colored in green, oxygen in red, nitrogen in blue and phosphorus in orange. Mg^2+^ is shown as a green ball.

**S6 Video Movie of a typical trajectory of the S2-GTP and the S2-open transition for Q61L.** The frames in these movies are rendered with the switch regions highlighted in green for S1 and blue for S2. The *α*3-helix is highlighted in yellow. The protein is shown as a ribbon with an transparent stick representation for the amino acids in S1 and S2. GTP is shown as solid sticks, with carbon atoms colored in green, oxygen in red, nitrogen in blue and phosphorus in orange. Mg^2+^ is shown as a green ball.

## References

1. Downward J. Targeting RAS signalling pathways in cancer therapy. Nature Reviews Cancer. 2003;3:11–22.

2. Lau KS, Haigis KM. Non-redundancy within the RAS oncogene family: Insights into mutational disparities in cancer. Molecules and Cells. 2009;28(4):315–320. doi: 10.1007/s10059-009-0143-7

3. Prior IA, Lewis PD, Mattos C. A Comprehensive Survey of Ras Mutations in Cancer. Cancer Research. 2012;72(10):2457–2467. doi: 10.1158/0008-5472.CAN-11-2612

4. Hobbs GA, Der CJ, Rossman KL. RAS isoforms and mutations in cancer at a glance. Journal of Cell Science. 2016;129(7):1287–1292. doi: 10.1242/jcs.182873

5. Dharmaiah S, Tran TH, Messing S, Agamasu C, Gillette WK, Yan W, et al. Structures of N-terminally processed KRAS provide insight into the role of N-acetylation. Scientific Reports. 2019;9(1). doi: 10.1038/s41598-019-46846-w

6. Pacold ME, Suire S, Perisic O, Lara-Gonzalez S, Davis CT, Walker EH, et al. Crystal Structure and Functional Analysis of Ras Binding to Its Effector Phosphoinositide-3-Kinase-*γ*. Cell. 2000;103:931–944.

7. Boriack-Sjodin PA, Margarit SM, Bar-Sagi D, Kuriyan J. The structural basis of the activation of Ras by Sos. Nature. 1998;394:337.

8. Spoerner M, Hermann C, Vetter IR, Kalblitzer HR, Wittinghofer A. Dynamic properties of the Ras switch I region and its importance for binding to effectors. Proc Natl Acad Sci USA. 2001;98:4944.

9. Kobayashi C, Saito S. Relation between the conformational heterogeneity and reaction cycle of Ras: molecular simulation of Ras. Biophys J. 2010;99:3726.

10. Spoerner M, Hozsa C, Poetzl JA, Reiss P K amd Ganser, Geyer M, Kalblitzer HR. Conformational states of human rat sarcoma (Ras) protein complexed with its natural ligand GTP and their role for effector interaction and GTP hydrolysis. J Biol Chem. 2010;285:39768.

11. Geyer M, Schweins T, Herrmann C, Prisner T, Wittinghofer A, Kalbitzer HR. Conformational Transitions in p21rasand in Its Complexes with the Effector Protein Raf-RBD and the GTPase Activating Protein GAP†. Biochemistry. 1996;35(32):10308–10320. doi: 10.1021/bi952858k

12. Hunter JC, Manadhar A, Carrasco MA, Gurbani D, Gondi S, Westover KD. Biochemical and Structural Analysis of Common Cancer-Associated KRAS Mutations. Mol Cancer Res. 2015;9:1325.

13. Buhrman G, Kumar VS, Cirit M, Haugh JM, Mattos C. Allosteric modulation of Ras-GTP is linked to signal transduction through RAF kinase. J Biol Chem. 2011;286:3323.

14. Fetics SK, Guterres H, Kearney BM, Buhrman G, Ma B, Nussinov R, et al. Allosteric effects of the oncogenic RasQ61L mutant on Raf-RBD. Structure. 2015;23:505–516.

15. Kyte J, Doolittle RF. A simple method for displaying the hydropathic character of a protein. Journal of Molecular Biology. 1982;157(1):105 – 132. doi: https://doi.org/10.1016/0022-2836(82)90515-0

16. Prakash P, Gorfe AA. Overview of simulation studies on the enzymatic activity and conformational dynamics of the GTPase Ras. Molecular Simulation. 2014;40(10-11):839–847. doi: 10.1080/08927022.2014.895000

17. Muraoka S, Shima F, Araki M, Inoue T, Yoshimoto A, Ijiri Y, et al. Crystal structures of the state 1 conformations of the GTP-bound H-Ras protein and its oncogenic G12V and Q61L mutants. FEBS Letters. 2012;586(12):1715–1718. doi: 10.1016/j.febslet.2012.04.058

18. Muraoka S, Shima F, Araki M, Inoue T, Yoshimoto A, Ijiri Y, et al. Crystal structure of H-Ras WT in complex with GppNHp (state 1); 2012. Available from: http://dx.doi.org/10.2210/pdb4efl/pdb.

19. Sharma N, Sonavane U, Joshi R. Probing the wild-type HRas activation mechanism using steered molecular dynamics, understanding the energy barrier and role of water in the activation. European Biophysics Journal. 2014;43(2):81–95. doi: 10.1007/s00249-014-0942-4

20. Dellago C, Bolhuis PG, Csajka FS, Chandler D. Transition path sampling and the calculation of rate constants. The Journal of Chemical Physics. 1998;108(5):1964–1977. doi: 10.1063/1.475562

21. Rogal J, Bolhuis PG. Multiple state transition path sampling. J Chem Phys. 2008;129(22):224107.

22. Webb B, Sali A. In: Comparative Protein Structure Modeling Using MODELLER. John Wiley & Sons, Inc.; 2002. p. 5.6.1–5.6.37. Available from: http://dx.doi.org/10.1002/cpbi.3.

23. Jorgensen WL, Chandrasekhar J, Madura JD, Impey RW, Klein ML. Comparison of simple potential functions for simulating liquid water. The Journal of Chemical Physics. 1983;79(2):926. doi: 10.1063/1.445869

24. Lindorff-Larsen K, Piana S, Palmo K, Maragakis P, Klepeis JL, Dror RO, et al. Improved side-chain torsion potentials for the Amber ff99SB protein force field. Proteins: Structure, Function, and Bioinformatics. 2010;78(8):1950–1958. doi: 10.1002/prot.22711

25. Meagher KL, Redman LT, Carlson HA. Development of polyphosphate parameters for use with the AMBER force field. Journal of Computational Chemistry. 2003;24(9):1016–1025. doi: 10.1002/jcc.10262

26. Essmann U, Perera L, Berkowitz ML, Darden T, Lee H, Pedersen LG. A smooth particle mesh Ewald method. The Journal of Chemical Physics. 1995;103(19):8577. doi: 10.1063/1.470117

27. Hess B, Kutzner C, van der Spoel D, Lindahl E. GROMACS 4: Algorithms for Highly Efficient, Load-Balanced, and Scalable Molecular Simulation. J Chem Theory Comput. 2008;4(3):435–447. doi: 10.1021/ct700301q

28. Bussi G, Donadio D, Parrinello M. Canonical sampling through velocity rescaling. The Journal of Chemical Physics. 2007;126(1):014101. doi: 10.1063/1.2408420

29. Parrinello M, Rahman A. Polymorphic transitions in single crystals: A new molecular dynamics method. J Appl Phys. 1981;52(12):7182. doi: 10.1063/1.328693

30. Hess B. P-LINCS: A Parallel Linear Constraint Solver for Molecular Simulation. Journal of Chemical Theory and Computation. 2008;4(1):116–122. doi: 10.1021/ct700200b

31. Eastman P, Swails J, Chodera JD, McGibbon RT, Zhao Y, Beauchamp KA, et al. OpenMM 7: Rapid development of high performance algorithms for molecular dynamics. PLOS Computational Biology. 2017;13(7):1–17. doi: 10.1371/journal.pcbi.1005659

32. Ryckaert JP, Ciccotti G, Berendsen HJC. Numerical integration of the cartesian equations of motion of a system with constraints: molecular dynamics of n-alkanes. Journal of Computational Physics. 1977;23(3):327 – 341. doi: https://doi.org/10.1016/0021-9991(77)90098-5

33. Sivak DA, Chodera JD, Crooks GE. Time Step Rescaling Recovers Continuous-Time Dynamical Properties for Discrete-Time Langevin Integration of Nonequilibrium Systems. The Journal of Physical Chemistry B. 2014;118(24):6466–6474. doi: 10.1021/jp411770f

34. Chodera J, Rizzi A, Naden L, Beauchamp K, Grinaway P, Fass J, et al. choderalab/openmmtools: 0.14.0 - Exact treatment of alchemical PME electrostatics, water cluster test system, optimizations; 2018. Available from: https://doi.org/10.5281/zenodo.1161149.

35. Åqvist J, Wennerström P, Nervall M, Bjelic S, Brandsdal BO. Molecular dynamics simulations of water and biomolecules with a Monte Carlo constant pressure algorithm. Chemical Physics Letters. 2004;384(4):288 – 294. doi: https://doi.org/10.1016/j.cplett.2003.12.039

36. Humphrey W, Dalke A, Schulten K. VMD – Visual Molecular Dynamics. Journal of Molecular Graphics. 1996;14:33–38.

37. Bolhuis PG, Chandler D, Dellago C, Geissler PL. TRANSITION PATH SAMPLING: Throwing Ropes Over Rough Mountain Passes, in the Dark. Annual Review of Physical Chemistry. 2002;53(1):291–318. doi: 10.1146/annurev.physchem.53.082301.113146

38. Swenson DWH, Prinz JH, Noe F, Chodera JD, Bolhuis PG. OpenPathSampling: A Python Framework for Path Sampling Simulations. 1. Basics. Journal of Chemical Theory and Computation. 2019;15(2):813–836. doi: 10.1021/acs.jctc.8b00626

39. Swenson DWH, Prinz JH, Noe F, Chodera JD, Bolhuis PG. OpenPathSampling: A Python Framework for Path Sampling Simulations. 2. Building and Customizing Path Ensembles and Sample Schemes. Journal of Chemical Theory and Computation. 2019;15(2):837–856. doi: 10.1021/acs.jctc.8b00627

40. Bolhuis PG. Transition path sampling on diffusive barriers. Journal of Physics: Condensed Matter. 2003;15(1):S113.

41. Hunter JD. Matplotlib: A 2D graphics environment. Computing In Science & Engineering. 2007;9(3):90–95. doi: 10.1109/MCSE.2007.55

42. Stelzl LS, Hummer G. Kinetics from Replica Exchange Molecular Dynamics Simulations. Journal of Chemical Theory and Computation. 2017;13(8):3927–3935. doi: 10.1021/acs.jctc.7b00372

43. McGibbon RT, Beauchamp KA, Harrigan MP, Klein C, Swails JM, Hernández CX, et al. MDTraj: A Modern Open Library for the Analysis of Molecular Dynamics Trajectories. Biophysical Journal. 2015;109(8):1528 – 1532. doi: 10.1016/j.bpj.2015.08.015

44. Bai L, Breen D. Calculating Center of Mass in an Unbounded 2D Environment. Journal of Graphics Tools. 2008;13(4):53–60. doi: 10.1080/2151237x.2008.10129266

45. Juraszek J, Vreede J, Bolhuis PG. Transition path sampling of protein conformational changes. Chemical Physics. 2012;396:30–44. doi: 10.1016/j.chemphys.2011.04.032

